# Revisiting the Structure/Function Relationships of H/ACA(-like) RNAs: A Unified Model for Euryarchaea and Crenarchaea

**DOI:** 10.1101/016246

**Authors:** Claire Toffano-Nioche, Daniel Gautheret, Fabrice Leclerc

## Abstract

A structural and functional classification of H/ACA and H/ACA-like motifs is proposed from the analysis of the H/ACA guide RNAs which have been identified previously in the genomes of Euryarchaea (Pyrococcus) and Crenarchaea (Pyrobaculum). A unified structure/function model is proposed based on the common structural determinants shared by H/ACA and H/ACA-like motifs in both Euryarchaea and Crenarchaea. Using a computational approach, structural and energetic rules for the guide-target RNA-RNA interactions are derived from structural and functional data on the H/ACA RNP particles. H/ACA(-like) motifs found in Pyrococcus are evaluated through the classification and their biological relevance is discussed. Extra-ribosomal targets found in both Pyrococcus and Pyrobaculum are presented as testable gene candidates which might support the hypothesis of a gene regulation mediated by H/ACA(-like) guide RNAs.

## Introduction

The H/ACA guide RNAs are part of a RNP machinery including several proteins (L7Ae, Cbf5, Nop10 and Gar1) which catalyzes the uridine-to-pseudouridine isomerization. Homologs of the snoRNA H/ACA found in Eukarya, they are found in a widespread number of Archaea. This family of small guide RNAs was detected both in Euryarchaea (*Archaeoglobus fulgidus* [1], *Haloferax volcanii* [2], *Pyrococcus abyssi* [3, 4, 5]), Crenarchaea (*Sulfolobus solfataricus* [6]) and Nanoarchaea (*Nanoarchaeum equitans* [7]). Computational screens for H/ACA RNAs and their potential targets in the archaeal genomes suggest they are present in a variable number of copies among all the archaeal phyla [8, 5]. The natural targets are ribosomal RNAs but other RNAs might be targeted with functional implications [9, 10]. In higher eukaryotes, snRNAs are additional targets [11]; the U2 snRNA is one particular target which is also modified in *S. cerevisiae* by an H/ACA guide RNA under particular conditions [12]. The artificial modification of mRNAs by the H/ACA guide RNP machinery was shown to suppress nonsense codons by targeting the uridine present as the first letter of stop codons [13]. On the other hand, snoRNAs have also been involved in unusual roles: in alternative splicing in the case of the C/D box guide RNA HBII-52 [14] or as one of the RNA biomarkers for non-small-cell lung cancer in the case of the H/ACA guide RNA snoRA42 [15].

In archaea, extensive structural and functional studies have focused on the H/ACA RNAs and RNPs from *P. abyssi* [16, 17, 4, 18, 19, 5, 20, 21] and *P. furiosus* [3, 22, 23, 24, 25, 26, 5, 27, 28, 29]. These studies describe the structure/function relationships for this RNA guide machinery associated with the RNA fold, the RNA:RNA interactions between the guide and its target(s) and the RNA:protein contacts (between L7Ae and the K-turn or K-loop motif, between Cbf5 and the ACA box, etc.). More recently, the discovery of ‘atypical’ H/ACA motifs in the *Pyrobaculum* genus has suggested an alternate way for this machinery to assemble and achieve its function [30]. These non-canonical H/ACA motifs, further designated as H/ACA-like motifs, are detected specifically in this genus of Crenarchaea and differ from the canonical H/ACA motifs in that the lower stem is truncated. They include two long free single-stranded regions at both 5’ and 3’ ends forming a pseudo-internal loop and the generic hairpin with an embedded K-turn or K-loop motif. In *Pyrobaculum,* two more canonical H/ACA motifs are also found (sR201 and sR202) among the ‘atypical’ H/ACA motifs. These two particular guide RNAs exhibit a lower stem (5 to 6 base pairs) although slightly shorter than that of the canonical H/ACA motifs found in *Pyrococcus* (8 to 9 base pairs) while the other eight guide RNAs are fully truncated with no lower stem. From a structural point of view, one may then consider that sR201 and sR202 do correspond to canonical H/ACA motifs. Actually, the first annotated ‘atypical’ H/ACA motif sR201 was identified previously as a canonical H/ACA motif in *P. aerophylum* [19, 31, 32].

In this work, the structure/function relationships of H/ACA RNAs are revisited from the structural and functional data on the canonical H/ACA guide RNAs of *Pyrococcus.* With 7 genes corresponding to 11 H/ACA RNA motifs in its genome, *Pyrococcus* is one of the archaeal genus with so many genes for this class of sRNA. As shown before [5], one given H/ACA guide RNA may potentially target several positions in 16S or 23S rRNAs and one given position may happen to be a potential target for two different H/ACA guide RNAs. However, there is only a unique pair between a target and its associated guide RNA, i.e. only a few of the potential targeted positions are actually modified (“true targets”) while others are not modified (“false targets”). Vice versa, a position potentially targeted by two guide RNAs is actually modified by only one H/ACA guide RNA (“productive”). Here, we propose a structural and functional classification that gives a rational for a “true target” and its associated “productive” H/ACA guide RNA or for a “false target” and its associated “non-productive” H/ACA guide RNA. The classification, based on the 2D folds of guide RNAs and the stability of the guide-target RNA:RNA duplexes, is intended to discriminate between these “productive”/“non-productive” H/ACA complexes; the “productive” H/ACA complexes are expected to lead to the pseudouridylation of some RNA target(s) (rRNA, tRNA or other RNA) while the non-productive ones may eventually play some alternative role.

The consistency of the classification is tested for all the archaea (*Archaeoglobus, Haloferax, Holoarcula, Sulfolobus, Aeropyrum,* etc.) where some H/ACA guide RNAs and their respective target(s) were characterized. The H/ACA-like motifs, found in *Pyrobaculum,* are also examined to determine whether the same classification may apply in spite of their structural specificities. The structural and functional relevance of the H/ACA-like motifs proposed recently in *P. abyssi* [33] is also evaluated according to the classfication.

## 1. Materials and Methods

### 1.1 RNA-seq Data

The full RNA-seq data on *P. abyssi* are available from the work published recently [33] and from its supplementary material(http://rna.igmors.u-psud.fr/suppl_data/Pyro).

### 1.2 Structural alignments and representations

Most of the sequences of the H/ACA motifs from *P. abyssi* are available in RFAM [34, 35]: Pab21 or snoR9 (RFAM ID: RF00065), Pab35 or HgcF (RFAM ID: RF00058), Pab40 or HgcG (RFAM ID: RF00064), Pab105 or HgcE (RFAM ID: RF00060), Pab19, Pa160, and Pab91 [5]. The STOCKHOLM alignments were edited to split the RNA genes with multiple motifs (Pab35: Pab35.1, Pab35.2, Pab35.3; Pab105: Pab105.1, Pab105.2; Pab40: Pab40.1, Pab40.2) into their corresponding unique motifs. All the individual H/ACA motifs were then merged to generate a full alignment. For the sake of graphical representation, the motifs with a large internal loop, with a single strand region exceeding 10 nucleotides (Pab40.2 and Pab19), were omitted. The 2D structure representations were generated from the STOCKHOLM alignments using the R2R program [36]. The structure representations of the guide-target RNA pairs are also generated using the R2R program. The corresponding pairing energies are calculated using the RNAsnoop utility [37] from the Vienna Package (version 2.0) [38]. In the analysis of the energy contributions to the duplex stability, we refer to the 5’ and 3’ duplex as defined in RNAsnoop.

In the case of the H/ACA-like motifs from *Pyrobaculum* and other new H/ACA(-like) motifs (from *Pyrococcus*), the alignments were obtained from the UCSC genome browser in MAF format [39, 40] and then converted into an aligned FASTA and STOCKHOLM formats. Finally, R2R [36] decorators were added to generate the 2D structure representations. The 2D representations of hybrid guide-target RNAs were generated using R2R with specific options and decorators; some detailed examples to reproduce this kind of representations are provided elsewhere [41]. The structural analysis of the H/ACA sRNAs was performed using X3DNA (version 2.1) [42] and Curves+[43].

### 1.3 Computational RNomics

A multi-step workflow is used to search for possible H/ACA-like motifs in the genome of *P. abyssi* [46, 47] (Fig. 1). The H/ACA-like motifs are identified using the descriptor-based approach implemented in RNAMotif [44]. Based on the 2D structure of the H/ACA(-like) motifs identified in *Pyrobaculum* [30], six families are defined to consider all possible candidates with different stem lengths and bulge positions (Supplementary data: Listings 1 to 12). The two canonical H/ACA motifs *Pae* sR201 & sR202 are defined using a unique descriptor. The following sRNAs: *Pae* sR204, *Pae* sR208, *Pae* sR209, *Pae* sR210 are also defined by a common descriptor with a bulge nucleotide one base pair downstream (3’ end) from the two GA pairs of the K-turn motif. Each of the other sRNAs (*Pae* sR203, *Pae* sR205, *Pae* sR206, *Pae* sR207) is defined by a unique descriptor. The substitution between K-turn and K-loop motifs is allowed: two descriptors are used for each of the six families of sRNAs. All the descriptors are provided (as text files) in the supplemental material.

**Figure 1.**
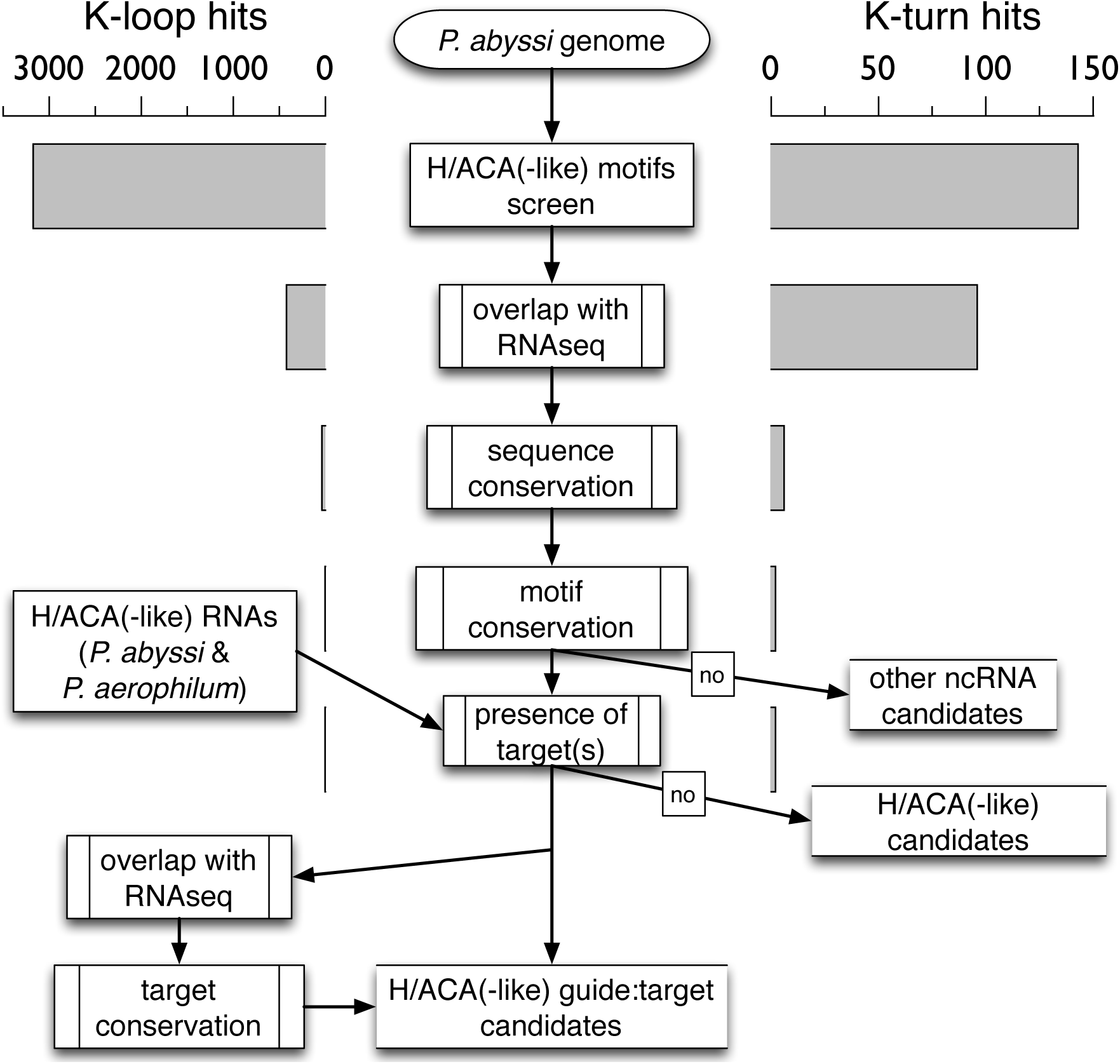
Workflow for the identification of H/ACA(-like) motifs and potential targets in *Pyrococcus abyssi* and *Pyrobaculum aerophilum.* The genome is initially screened for the presence of H/ACA-like motifs using a descriptor-based approach (RNAMotif [44]). The number of hits is given at each step of the workflow as an indicative value for the motifs carrying either a K-turn or K-loop submotif. The first filter is the selection of motifs in the non-coding transcriptome (S-MART scripts [33, 45]), then the selection of conserved sequences and motifs among other archaea (UCSC archaea genome browser [40, 39]) and finally the selection of motifs that have potential target(s) in the genome which are themselves transcribed and conserved among other archaea.

A series of S-MART scripts [45] is used to extract the H/ACA(-like) motifs that lie in the non-coding transcriptome of *P. abyssi.* A minimum overlap of 50 nucleotides is required to filter out the initial hits. The sequence and motif conservation are verified using the UCSC archaeal genome browser [39, 40]. A combination of two softwares: locaRNA [48] and RNAalifold [49] are used to identify optimal and sub-optimal 2D structures which are consistent with the H/ACA(-like) fold. The search for associated targets is performed using the RIsearch program [50], using the recommended extension penalty (“-d 30”). The potential RNA targets are identified by searching through the whole genomes matching sequences that can associate with a pseudo-guide sequence containing the 5’ and 3’ duplex elements from the internal loop of the guide RNA. This pseudo-guide sequence is obtained by merging the 5’ and 3’ duplex elements from the internal loop of the guide RNA separated by a di-nucleotide spacer equivalent to the di-nucleotide spacer containing the targeted U position. In order to impose a 5’-UN-3’ sequence constraint in the target RNAs, the di-nucleotide spacers in the pseudo-guide sequences are defined by the complementary element: 5’-NA-3’. The target RNA candidates are selected based upon a RIsearch energy score of −16 kcal/mol or lower which corresponds to the less favorable energy score obtained for validated guide-pair targets (with a deviation of 1 kcal/mol). Only target sequences with at most two bulge nucleotides are tolerated but no bulge nucleotide is allowed on the 5’ duplex (5’ end of the paired target sequence) upstream from the predicted modified position.

After a guide-target pair is identified, two additional filters are applied: the presence of the target sequence in a region of the genome of *P. abyssi* which is expressed [33], the conservation of the target in both *P. abyssi* and *P. aerophilum* whether it is present in homologous genes (identified using the BBH approach [51]) or in genes which are functionally related. First, a functional annotation analysis using the DAVID bioinformatics resources [52, 53] is carried out to evaluate the enrichment in genes and family of genes targeted in both genomes. Then, the targeted genes that belong to the same functional category are analyzed and characterized using the tools described above for the representations and energy calculations of the guide-target pairs.

## 2. Results and Discussion

### 2.1 Current Structure/Function Model of H/ACA motifs

The H/ACA motif is well described as a stem-loop-stem motif closed by an apical loop and terminated by an ACA box at the 3’ end [54]. In archaea, the L7Ae ribosomal protein is part of the H/ACA RNP particle and specifically binds to a K-turn motif which is embedded in the upper stem or merged within the apical loop as a K-loop. The lower stem generally includes 9 base-pairs in the canonical H/ACA motifs and the distance between the ACA box and the base-pair closing the internal loop in the upper stem is around 15 nt (between 14 and 16 nt [1]); in the numerous H/ACA motifs present in *P. abyssi,* an empirical constraint between 14 and 15 nt was proposed [5]. On the other hand, no well-defined constraint is proposed regarding the position of the K-turn or K-loop motif with respect to the internal loop. In *Haloferax volcanii,* the distance between the base-pair closing the internal loop in the upper stem and the second sheared G:A base-pair of the K-turn or K-loop motif corresponds to 8 base-pairs [2]. In the same species, the 2D folds proposed for the “productive” H/ACA guide RNAs involve a distance of 9 base-pairs [1, 16, 55] or 10 base-pairs [1, 16]. In *Pyrococcus,* it is between 9 and 11 base-pairs but it is restrained to 9 or 10 base-pairs in the “productive” H/ACA guide RNAs [5]. Finally, in the H/ACA-like motifs discovered in *Pyrobaculum,* this constraint can vary between 9 and 12 base-pairs [30]. When searching for new H/ACA(-like) motifs in the archaeal genomes, a very loose constraint is generally used for this distance: between 5 and 12 nucleotides [55, 5, 8, 19]. In all the 3D structures of the full H/ACA RNP complexes (L7Ae, Nop10, Cbf5, and Gar1) including or not a target RNA (PDB IDs: 2HVY, 3LWO, 3HAX) [23, 27, 28], this distance constraint is exactly 10 base-pairs: it gives the proper position and orientation of the K-turn or K-loop motif within the RNP complex for L7Ae to bind the guide RNA and interact with Nop10. The only exception is the case of the incomplete H/ACA RNP complex reconstituted with a composite guide RNA derived from Pf9 (Pa160 ortholog) which exhibits a 9 base-pairs constraint (PDB ID: 2RFK) [26]. Interestingly, the L7Ae protein is missing in this particular H/ACA RNP complex. To determine whether these constraints apply to all the known “productive” H/ACA guide RNAs, we are revisiting the structure/function relationships of the archaeal H/ACA and H/ACA-like motifs.

### 2.2 New Structural and Functional Classification of H/ACA RNAs in Pyrococcus

The sequence and structure analysis of H/ACA guide RNAs from *P. abyssi* reveals the existence of different structural families which can all fit into a common structure/function model consistent with a so-called ‘10/n/9’ fold (Fig. 2) where the numbers indicate a distance in base-pairs: (a) from the internal loop to the K-turn or K-loop motif and (b) from the internal loop to the ANA box (which is generally equivalent to the length of the lower stem), respectively. The 2D consensus structure (Fig. 2(a)A), represented from a STOCKHOLM alignment (Supplementary Listing 13) is consistent with the 3D structure determined for a chimeral H/ACA RNA from *P. furiosus* (Fig. 2(a)B, PDB ID: 3LWO) [28] (patched from the Pf9 and Pf6 RNAs from *P. furiosus* that are orthologs of Pa160 and Pa35.2 from *P. abyssi*). The main features of the consensus (Fig. 2(a)A) are the following: a lower stem with 9 base pairs, a symmetric or asymmetric internal loop (usually small: 5-8 nt and 5-6 nt on the 5’ and 3’ single-stranded region, respectively), an upper stem including 10 stacked base-pairs between the internal loop and the K-turn/K-loop motif.

**Figure 2.**
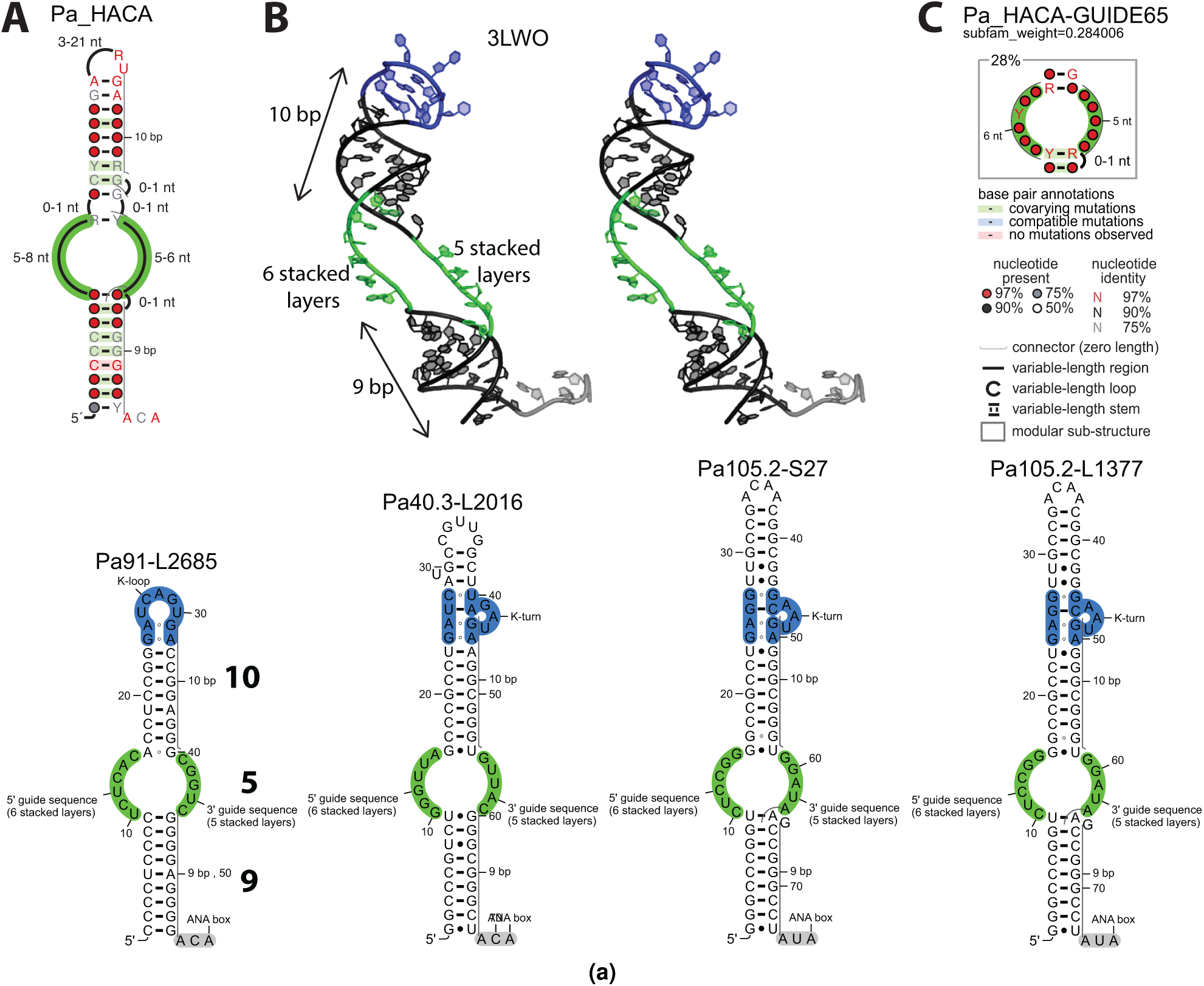
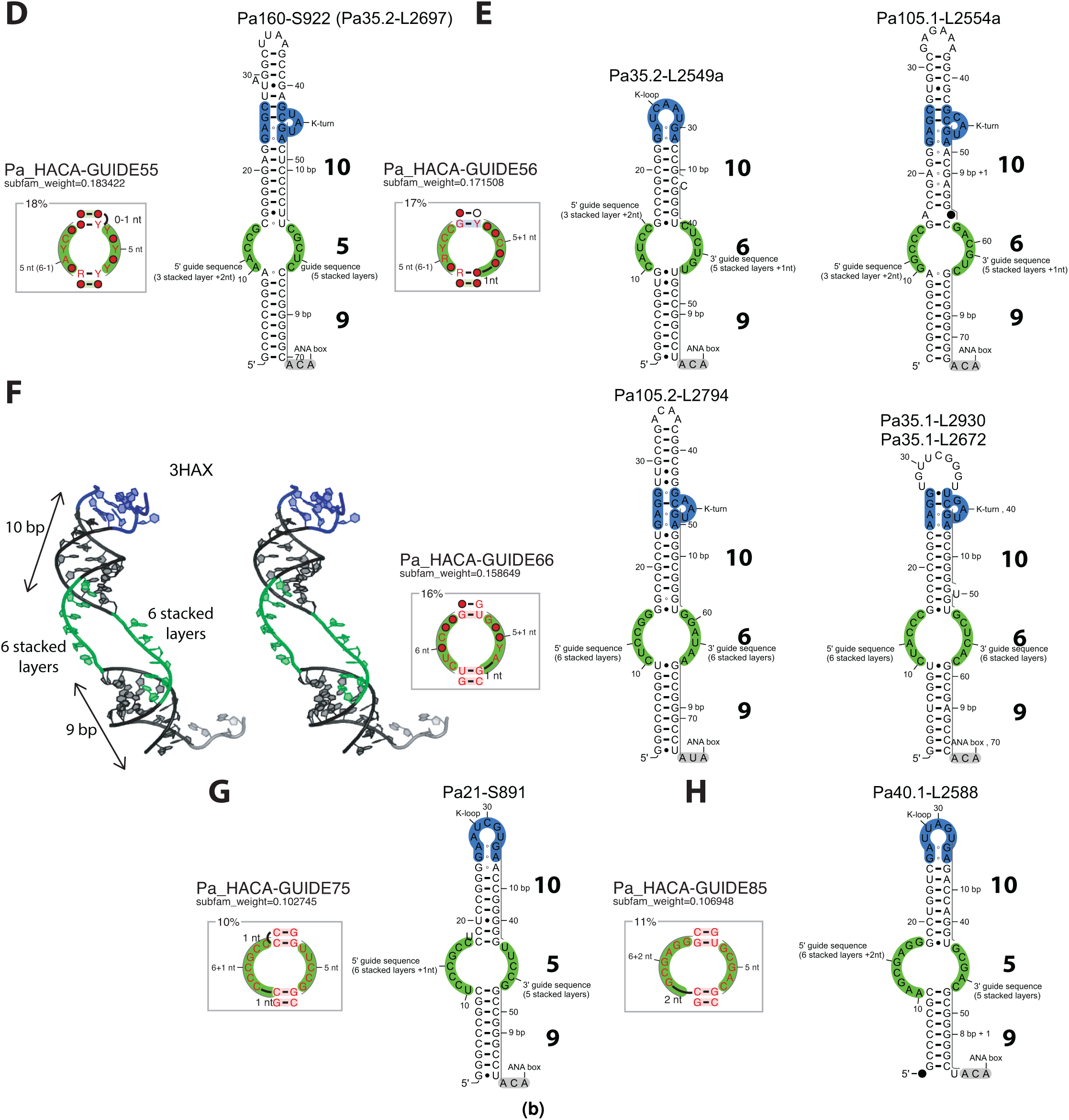
Structure/Function Model of H/ACA guide RNAs in *Pyrococcus abyssi.* **A.** The consensus 2D structure, labeled Pa HACA, is obtained from a curated STOCKHOLM alignment modified from the RFAM entries for H/ACA guide RNAs in P. abyssi (RFAM [34, 35] IDs: RF00058, RF00060, RF00065). **B.** The 3D structure of the chimeric guide RNA from *Pyrococcus furiosus* is shown in a stereo view as it is in the H/ACA sRNP complex including its RNA target (PDB ID: 3LWO) [28]. **C.** The more representative structural subfamily Pa HACA-GUIDE65 (6 and 5 nucleotides in the 5’ and 3’ strand regions of the internal loop, respectively) belongs to the ‘10/5/9’ fold family. The corresponding H/ACA guide RNAs from this subfamily in *P. abyssi* are shown: Pa91, Pa40.3, Pa105.2. Each RNA fold is annotated with the names of the H/ACA motif and its associated target in the large (L) or small (S) ribosomal subunit indicated by the position which is modified in the rRNAs. The graphical representations of the full 2D structures and the internal loop subfamilies were generated using the R2R program [36]. The internal loop of the H/ACA motif is shaded in green, the K-turn or K-loop motifs in blue and the ANA motif in grey. **D.** Structural subfamily Pa HACA-GUIDE55 and one of its representative folds (Pa160-S922; the Pab35.2-L2697 fold is equivalent). **E.** Structural subfamily Pa HACA-GUIDE56 and its 2 members (Pa35.2-L2549 and Pa105.1-L2554). **F.** Structural subfamily Pa HACAGUIDE66 corresponding to the 3D structure of the chimeric guide RNA derived from Afu46 (from *Archaeoglobus fulgidus* and its representative folds (Pa105.2-L2794 and Pa35.1-L2930 or Pab35.1-L2672). Only the RNA component of H/ACA RNP complex (PDB ID: 3HAX) is shown in a stereoview. **G.** Structural subfamily Pa HACA-GUIDE75 and its representative fold (Pa21-S891). **H.** Structural subfamily Pa HACA-GUIDE85 and its representative fold (Pa40.1-L2588). Pa35.2 can adopt two different folds: 10/6/9 or 10/5/9 whether the base-pair from the apical stem closing the internal loop is a G:U (Pab35.2-L2549a) or G:C (Pa35.2-L2697), respectively.

The H/ACA motifs are then classified in two fold families denoted ‘10/5/9’ (Fig. 2(a)-(b)) and ‘10/6/9’ (Fig. 2(b)) where n={5,6} indicates the size of the 3’ single-stranded region of the internal loop. The ‘10/5/9’ fold family is consistent with the 3D structure of a chimeral H/ACA RNA from *P. furiosus* (Fig. 2(a)B) while the ‘10/6/9’ fold family is consistent with the 3D structure of another chimeral H/ACA RNA (Fig. 2(b)F) derived from *Archaeoglobus fulgidus* (Afu46) which is assembled with the H/ACA ribonucleoproteins from *P. furiosus* (PDB ID: 3HAX) [23]. The sequence of Afu46 is somehow similar to that of Pf91 in *P. furiosus* [5] (homolog of the Pa91 motif (Fig. 2(a)) from *P. abyssi*). In the ‘10/6/9’ fold family, the internal loop is extended by the addition of one nucleotide in the 3’ single-stranded region of the internal loop which folds with 6 stacked nucleotides (Fig. 2(b)E-F). As a result, the number of stacked layers from the ANA box to the closing base-pair of the upper stem differs in the two fold families ‘10/5/9’ and ‘10/6/9’ with 14 and 15 nucleotides, respectively. Nevertheless, the two crystallized H/ACA guide RNAs (PDB IDs: 3LWO and 3HAX, Fig. 2(a)B and Fig. 2(b)F) which are representative from both fold families do exhibit the same relative positions of the internal loop, lower stem and ANA box when superposing the two RNP complexes. In the ‘10/6/9’ family, the additional stacked nucleotide in the 3’ end duplex sits at the location of the first base-pair from the lower stem in the ‘10/5/9’ family. This is made possible by a change in the morphology of the lower stem where the RNA helix is more bent in ‘10/6/9’. This more pronounced bending is associated with a more compact helix (smaller helical rise) by almost 1Å and less twisted (smaller helical twist) by about 15 degrees (Table S1). At the junction between the lower stem and the internal loop, the helical twist is large in ‘10/5/9’ (43.1°) but small in ‘10/6/9’ (30.8°) with respect to a canonical A-RNA helix (33.6°). As suggested in a previous work [21], the RNA sequence may have an influence on the intrinsic bending of the lower stem of H/ACA guide RNAs. It is likely that the proteins of the RNP particle also play a non-specific role to induce a bending suitable with its catalytic activity in this portion of the guide RNA.

Different structural subfamilies can then be distinguished based on the size of the 5’ single-stranded region of the internal loop, e.g. the Pa HACA-GUIDE65 subfamily corresponds to an asymmetric internal loop with 6 and 5 residues, respectively (Fig. 2(a)C). The 2D representation of each guide RNA corresponds to the expected “productive” fold in the H/ACA RNP particle and thus includes the name of the H/ACA motif (e.g. Pa91) and its associated target in the large (e.g. L2685) or small (e.g. S27) ribosomal subunit. In total, 6 different subfamilies are present and the Pa HACA-GUIDE65 subfamily is the larger one (28%) including 4 H/ACA guide RNAs in *Pyrococcus* (Fig. 2(a)). The other subfamilies, represented in the order of the frequence of occurence in all H/ACA motifs (Fig. 2(b)D-H), are variations from the Pa HACA-GUIDE65 subfamily in the size of the internal loop which is either shortened or extended by one or two nucleotides on either sides. H/ACA RNA homologs in other species can switch to a different subfamily due to the insertion(s) or deletion(s) in the internal loop. In the Pa HACA-GUIDE55 family, Pa160 has one homolog in *P. furiosus:* Pf9 that switches to the Pa HACA-GUIDE65 family. The homologs of Pa35.2 in *Thermococcus kodakaraensis* and Pa105.2 in *P. furiosus* switch from Pa HACA-GUIDE56 to Pa HACA-GUIDE66. The homolog of Pa40.1 in *T. kodakaraensis* switches from Pa HACAGUIDE85 to Pa HACA-GUIDE65 (data not shown).

Some other minor structural variations are observed (Fig. 2). The first one corresponds to the loss of the first base-pair at the bottom of the lower stem in Pa40.1 (Pa40.1-L2588 in Fig. 2(b)H). However, the distance between the first base-pair of the lower stem and the ANA box remains the same: 9 nucleotides (8 base-pairs and one unpaired U nucleotide). In other closely related species like *P. horikoshii, P. yayanossi,* and *T. kodakaraensis,* the first base-pair is preserved as a wobble U-G or U-A base-pair. The second minor variation involves a bulge nucleotide in Pa105.1 (A at position 16: Pa105.1-L2554a in Fig. 2(b)E) which is assumed to be stacked in the upper stem: this conformation is supported by compensatory changes in the ortholog from *T. kodakaraensis:* the substitution of the bulge nucleotide by a base-pair which is correlated with the substitution of the two unusual G-A sheared base-pairs embedded in the upper stem by two canonical base-pairs (Supplementary Fig. S1). The presence of both a single bulge nucleotide and two G-A base-pairs in the upper stem are expected to allow the adjustment of the helical twist so that the relative position of the K-turn/K-loop motif with respect to the internal loop is preserved (a bulge A nucleotide adopts such a stacked conformation in the 24-mer RNA hairpin coat protein binding site from bacteriophage R17, PDB ID: 1RHT [56]).

From this classification, we can summarize the structural determinants of H/ACA RNAs that reside in the relative positions of both the K-turn/K-loop motif and the ANA box with respect to the internal loop. The distance between the K-turn/K-loop motif and the internal loop includes 10 stacked layers. A more detailed analysis also suggests a prevalence of the 5’ duplex over the 3’ duplex in the guide-target hybrid duplex. The “productive” H/ACA RNA are thus expected to fulfill the following criteria: (1) a first stretch of 10 stacked layers from the upper stem to the K-turn/K-loop motif, (2) a second stretch of 14 to 15 stacked layers from the lower stem to the ANA box. The second stretch is generally composed of 9 base-pairs in the lower stem and 5 to 6 nucleotides in the 3’ region of the internal loop. In addition, the 5’ region of the internal loop should not have fewer base-pairs than the 3’ region in the RNA guide-target hybrid duplex due to geometrical constraints. All the H/ACA guide RNAs that were validated from a functional point of view in *P. abyssi* [5] fit into one of the two fold families: ‘10/6/9’ or ‘10/5/9’ (Fig. 2), except Pa19 for which an atypical pairing model was proposed with only 8 base-pairs in the upper stem (‘8/7/9’ fold); this particular case is discussed later. Pa105.1-L2554 (Fig. 2(b)) was initially considered as a ‘9/6/9’ fold but the structural and phylogenetic data suggest it actually fits into the ‘10/6/9’ fold (Fig. 2(b)E and Supplementary Fig. S1).

Because of the conformational flexibility of RNA, the criteria relative to the size of the internal loop are only relevant to the active conformation of the H/ACA RNAs while they are associated to the proteins of the H/ACA RNP particle and the RNA target. Therefore, one can imagine the internal loop may change in size due to the opening of the lower or upper stems at the closing base-pairs to accommodate different RNA targets (Fig. 3). The opening of the upper stem systematically leads (if not compensated) to switch from a ‘productive’ (’10/5(6)/9’) to a ‘non-productive’ fold (‘9/5(6)/9’, ‘8/5(6)/9’, etc.). The Pa35.1 motif is productive (‘10/6/9’) with the following ribosomal targets: L2930 and L2672 (23S rRNA) but non-productive with other targets in the 16S rRNA: S922 (‘9/7/9’) or S1122 (‘9/8/9’) (Figure 3(a)A). On the opposite, the opening of the lower stem can preserve the 14 or 15 stacked nucleotides allowing extended single-stranded regions for base-pairing with RNA target(s). One example is provided by Pa35.2-L2549 (‘10/6/9’) where the lower stem is shortened by the opening of one (‘10/7/8’) or eventually two wobble base-pairs (‘10/8/7’).

**Figure 3.**
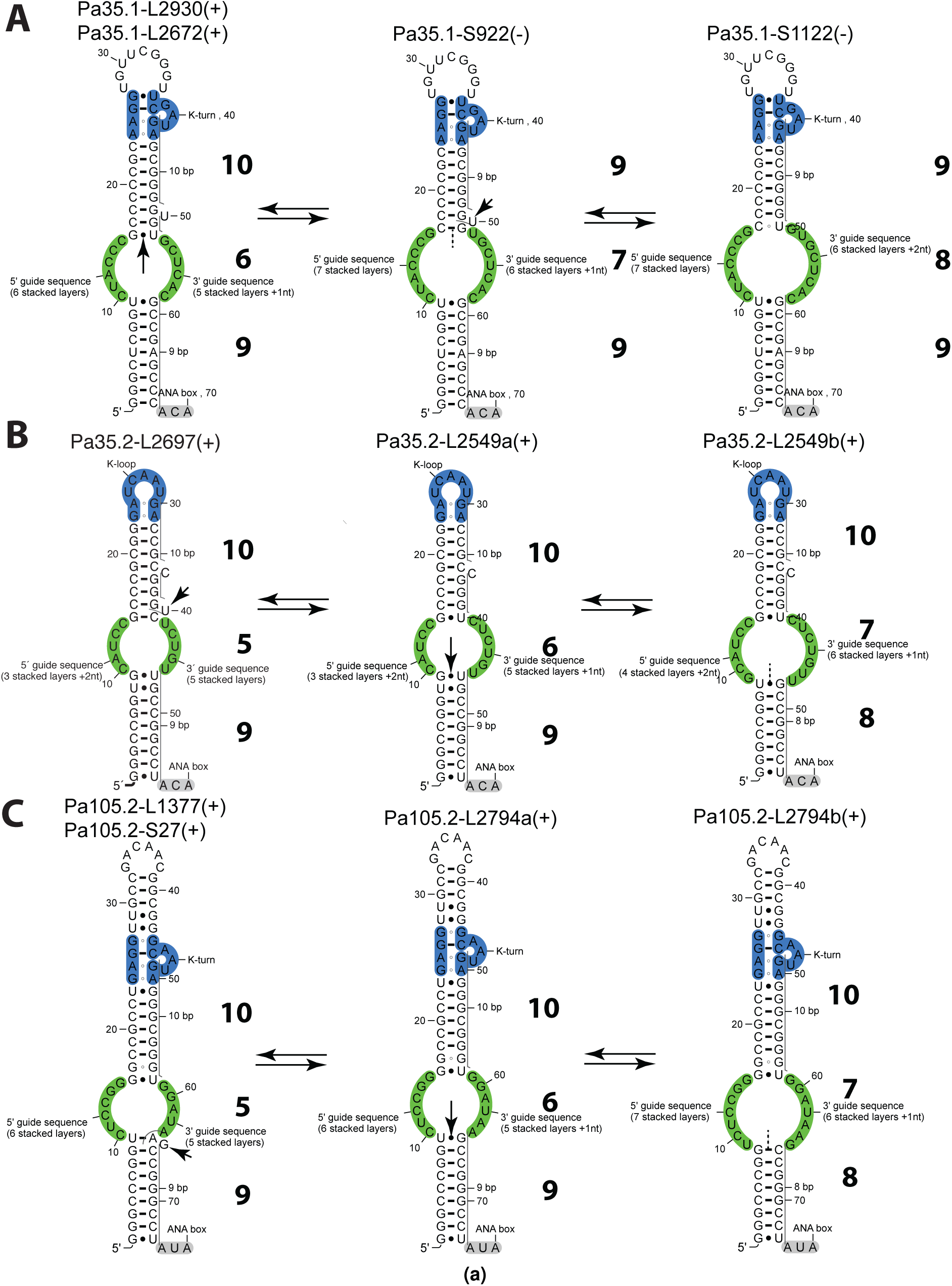
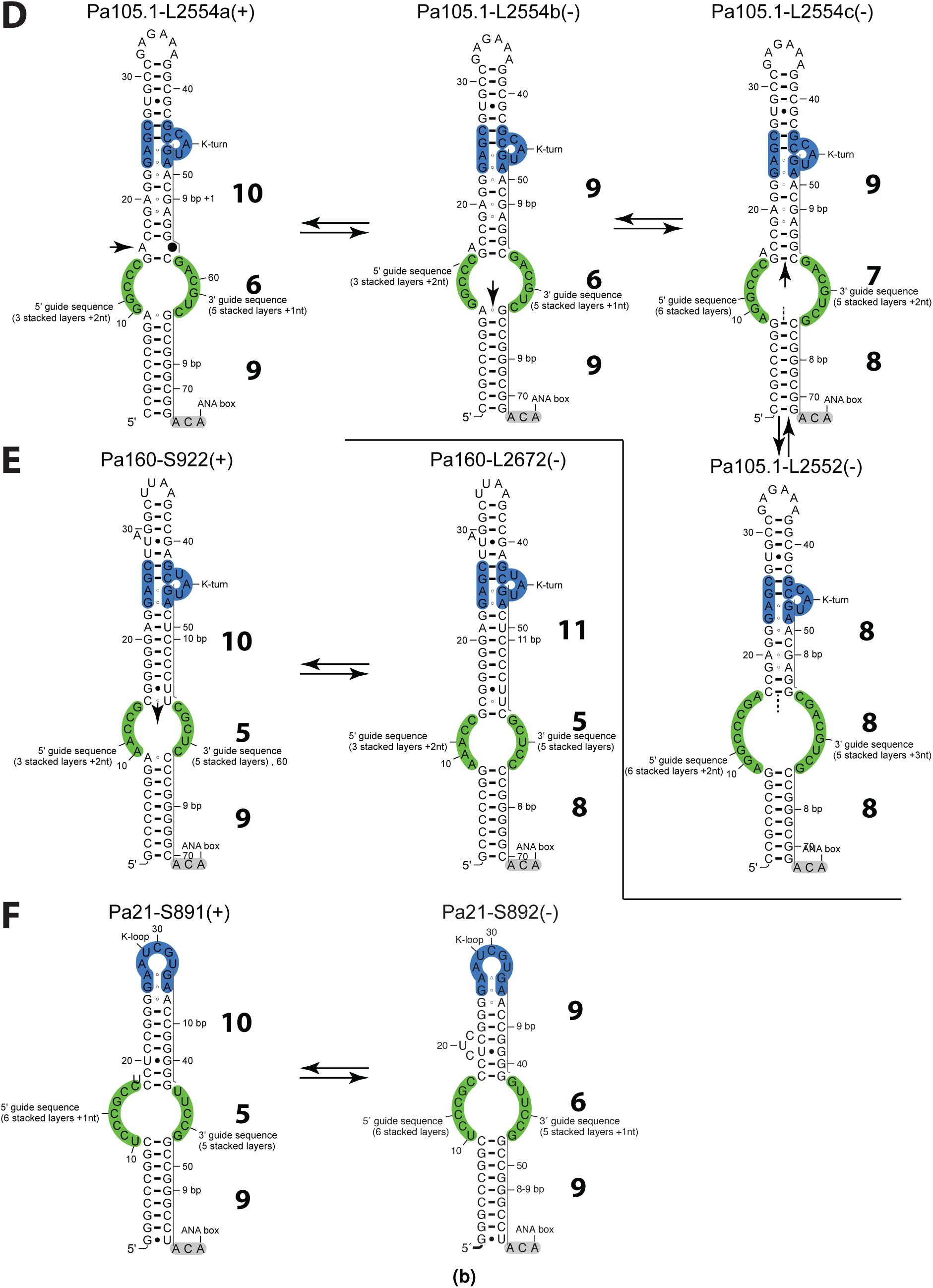
Productive/Non-productive H/ACA guide RNAs. **A.** Productive/non-productive folds for Pa35.1 and its true (L2930, L2672) and false (S922 and S1122) targets. **B.** Productive folds for Pa35.2 and its true targets: L2697 and L2549. **C.** Productive folds for Pa105.2 and its true targets: L1377, S27, L2794. The 2D structures of the guide RNAs are shown according to the RNA fold consistent with the pairing with its associated target (the RNA targets are omitted for clarity): productive and non-productive complexes are indicated by a (+) and (-) sign, respectively. The different possible structures for a given guide RNA are shown using a double arrow. Structural changes associated with opening or closing base-pairs in the internal loop are indicated by single vertical arrows; dashed lines indicate the opening of the internal loop. Bulge nucleotides switching out/in of a stem are marked using an oblique arrow. **D.** Productive/non-productive folds for Pa105.1 ands its true targets: L2554, false target: L2552. **E.** Productive/non-productive folds for Pa160 ants its true (S922) and false (L2672) targets. **F.** Productive/non-productive folds for Pa21 and its true (S891) and false (S892) targets. Pa105.1 can adopt alternative folds by opening the lower stem by one base-pair to increase the base-pairing between the guide and its target (x/6/9 to x/7/8). Opening the apical stem as well by one base-pair (x/8/8) creates an internal loop that would accept L2552 as possible target. Adding one base-pair at the bottom of the lower stem of Pa160 creates an internal loop that would accept L2672 as possible target.

The presence of a bulge nucleotide in some specific positions near the base pairs closing the internal loop can help switching between different “productive” folds, thus extending the repertoire of potential targets. Bulge nucleotides are a source of sequence variability in the internal loop when they are located either at the (n-1) position from the closing base-pair of the upper stem (e.g. Pa35.2-L2697, Fig. 3(a)B) or at the (n+1) position from the closing base-pair of the lower stem (e.g. Pa105.2-L1377 or Pa105.2-S27, Fig. 3(a)C). The 5’ region of the internal loop is thus enlarged by the insertion of one nucleotide at the 5’ or 3’ end while preserving a canonical base-pair (Watson-Crick or wobble) closing the internal loop. In the first case, the sequence is modified at the position corresponding to the last base-pair in the 5’ duplex of the guide-target hybrid. In the second case, the sequence is extended at the first position of the 5’ duplex. In the case of Pa35.2- L2697, the closing base-pair G15-C41 imposes a ‘10/5/9’ fold; the bulge nucleotide U-40 can form a wobble pair with G15 by pushing C-41 into the internal loop and thus induce a switch to the ‘10/6/9’ fold which is productive in Pa35.2- L2549 (Fig. 3(a)B). Similarly, the Pa105.2 motif can switch from a ‘10/5/9’ (Pa105.2-L1377 or P105.2-S27)subfamily to a ‘10/6/9’ fold by substituting the closing base pair from the lower stem: U9-A64 by the U9-G65 while pushing A64 in the internal loop (Pa105.2-L2794). The Pa105.2 motif can also undergo typical rearrangements described above for the opening of the lower stem: from ‘10/6/9’ to ‘10/7/8’ or alternatively from ‘10/5/9’ to ‘10/6/8’ (Fig. 3(a)C).

In the case of Pa105.1, various examples of possible changes that impact negatively the upper stem are also shown (Fig. 3(b)D). Pa160-S922 (‘10/5/9’) is productive while Pa160- L2672 (‘11/5/8’) obtained by ‘zipping down’ the internal loop is not productive (Fig. 3(b)E). More significant conformational changes associated with a fold change are possible for a few H/ACA motifs; Pa21-S891 (‘10/5/9’) is productive while its alternate fold Pa21-S892 (‘9/6/9’) is not (Fig. 3(b)F). Two particular motifs have potential crossed targets: Pa160 and Pa35.1 which can both target the positions S922 and L2672 (Fig. 3(a)-(b)). The productive complexes are Pa160-S922 (‘10/5/9’) and Pa35.1-L2672 (‘10/5/9’) while the nonproductive ones are: Pa35.1-S922 (‘9/7/9’) and Pa160-L2672 (‘11/5/8’). All the productive complexes do exhibit 10 base-pairs or stacked layers between the the upper stem and the K-turn or K-loop motif. A bulge in the upper stem may compensate the presence of only 9 base-pairs if it is stacked in the helix (and most likely stabilized in the RNP complex). The productive complexes are also made up of 14 or 15 stacked layers from the closing base-pair of the upper stem to the ANA box including 7 to 9 base-pairs in the lower stem depending on the extent of opening of the internal loop.

The structure-function classification based on the structure and pairing models inferred from the available data discriminates the ‘productive’ from the ‘non-productive’ H/ACA guide RNAs for all possible guide-target pairs described previously taking into account one additional criterion for the guide-target association. The calculated stability of the guide-target complex (RNAsnoop [37]) is not discriminative enough since some non-productive complexes are as stable as productive ones (compare for example Pa21-S891 and Pa21-S892 in Table 1). However, a cutoff value can be defined for the prediction of new guide-pair targets as the minimum energy value for a productive complex which is −27 kcal/mol for Pa35.2- L2697 (Table 1). According to the classification, 17 guide-target pairs correspond to ‘productive’ folds: e.g. ‘10/6/9’, ‘10/5/9’ or equivalent folds (see Table 1), while 8 other pairs correspond ‘non-productive’ folds. All the guide RNAs which are predicted as ‘non-productive’ are indeed non-productive (6 true negative cases). All the guide RNAs which are productive are also predicted as ‘productive’ (15 true positive cases). The two missing cases (false positives) correspond to guide-target pairs which adopt a ‘productive’ fold with a weak interaction at 5’ duplex: Pa35.2-L2250 and Pa105.2- S995 (Table 1). This duplex includes only 3 base-pairs, two of which are wobble base-pairs for Pab35.2-L2250 (Supplementary Fig. S2) or 4 base-pairs, two of which are also wobble base-pairs for Pa105.2-S995. The only major difference between the two guide-target complexes Pa105.2-L1377(+) (Fig. 4A and Supplementary Listing 14) and Pa105.2-S995(-) (Fig. 4B and Supplementary Listing 15) is the pairing in the 5’ duplex of Pa105.2 (upstream from the pseudo-uridine position): a perfect match for the productive complex but a poor match for the non-productive complex. In the other target in 23S rRNA (L2794), the 5’ duplex energy is more stable (−8.5 kcal/mol) for the productive complex Pa105.2- L2794 (Fig. 4C and Supplementary Listing 16). Although the total energy of both guide-RNA complexes is the same, the 5’ duplex energy is −5.7 kcal/mol for the non-productive complex (Pa105.2-S995) and -6.9 kcal/mol for the productive one (Pa105.2-L1377). Similarly, the 5’ duplex energy of Pab35.2-L2250 is only −5.8 kcal/mol. In some extreme case, the 3’ duplex energy of Pab35.2-L2549(+) is very weak (−1 kcal/mol) but the 5’ duplex energy is still below −6 kcal/mol (−9.5 kcal/mol). We can expect the existence of a thermodynamic control where a minimum stability of the 5’ duplex is required for the guide-target complex to be productive. This rule is followed in all negative cases. In the border line case of Pa40.1-L2588, the 5’ duplex energy is −6.2 kcal/mol (Table 1). A minimum energy of −6 kcal/mol can be used as a discriminant criterion to exclude guide-target pairs which may not be productive (Table 1).

**Table 1.** Functional and structural features from known H/ACA guide RNAs in *Pyrococcus* targeting 16S and 23S rRNAs.

| guide-target pairs | conservation<sup>a</sup> | expression<sup>b</sup> | promoter<sup>b</sup> | % GC content | pseudouridylation<sup>c</sup> | structure-function model<sup>d</sup> | hybrid energy<sup>e</sup> | duplex energy<sup>f</sup> | 5' duplex energy<sup>f</sup> |
| --- | --- | --- | --- | --- | --- | --- | --- | --- | --- |
| Pa19-S1017<sup>g</sup> | + | + | - | 72 | + | 8/7/9- | -38 | -24 | -11 |
| Pa19-S1017<sup>h</sup> | + | + | - | 72 | + | 10/6/9+ | -38 | -21 | -11 |
| Pa21-S891 | + | + | + | 76 | + | 10/5/9+ | -29 | -20 | -7.7 |
| Pa21-S892 | + | + | + | 76 | - | 9/6/9- | -29 | -21 | -7.7 |
| Pa35.1-L2930 | + | + | + | 70 | + | 10/6/9+ | -30 | -19 | -8.1 |
| Pa35.1-L2672 | + | + | + | 70 | + | 10/8/7+ | -30 | -17 | -16 |
| Pa35.1-S922 | + | + | + | 70 | - | 9/7/9- | -28 | -13 | -6.5 |
| Pa35.1-S1122 | + | + | + | 70 | - | 9/8/9- | -28 | -15 | -9.4 |
| Pa35.2-L2549 | + | + | + | 71 | + | 10/6/9+ | -29 | -6.4 | -9.5 |
| Pa35.2-L2697 | + | + | + | 71 | + | 10/5/9+ | -27 | -10 | -6.0 |
| Pa35.2-L2250 | + | + | + | 72 | - | 10/5/9+ | -27 | -6.8 | -5.1 |
| Pa40.1-L2588 | + | + | - | 69 | + | 10/5/9+ | -30 | -21 | -6.2 |
| Pa40.2-L1932 | + | + | - | 74 | + | 10/6/8+ | -46 | -13 | -11 |
| Pa40.2-L278 | + | + | - | 74 | - | 9/7/8- | -46 | -20 | -13 |
| Pa40.3-L2016 | + | + | - | 63 | + | 10/5/9+ | -32 | -13 | -7.7 |
| Pa91-L2685 | + | + | + | 72 | + | 10/5/9+ | -33 | -18 | -11 |
| Pa105.1-L2554 | + | + | + | 76 | + | 10/6/9+ | -35 | -15 | -8.4 |
| Pa105.1-L2552 | + | + | + | 76 | - | 8/7/9- | -34 | -20 | -8.1 |
| Pa105.2-S995 | + | + | + | 71 | - | 10/6/9+ | -34 | -13 | -5.7 |
| Pa105.2-L2794 | + | + | + | 71 | + | 10/6/9+ | -34 | -15 | -8.5 |
| Pa105.2-L1377 | + | + | + | 71 | + | 10/6/9+ | -34 | -13 | -6.9 |
| Pa105.2-S27 | + | + | + | 71 | + | 10/6/9+ | -34 | -17 | -6.9 |
| Pa160-L2672 | + | + | + | 69 | - | 11/5/8- | -38 | -12 | -8.7 |
| Pa160-S922 | + | + | + | 69 | + | 10/5/9+ | -38 | -20 | -14 |

<sup>a</sup> as provided by the UCSC genome browser [39, 40]. <sup>b</sup> as determined from RNA-seq data [33]. <sup>c</sup> as determined from previous work by RT-CMCT analysis and *in vitro* activity of pseudouridylation [5]. <sup>d</sup> structure-function model as proposed from the 'productive'/'non-productive' classification (the +/- sign indicates whether it is predicted to be productive or not); the listed models also include some additional models that optimize the hybrid and duplex energies: 10/6/8 (Pa40.2-L1932) and 10/8/7 (Pa35.1-L2672) which are derived from 10/5/9 and 10/6/9 by opening the lower stem by one or two base-pairs (Fig. 3), respectively. <sup>e</sup> as calculated by RNAsnoop (kcal/mol) from the Vienna RNA package [37]. <sup>f</sup> the duplex energy includes both the 5' and 3' duplex energies with a correction factor (+4.1 kcal/mol). <sup>g</sup> the Pa19-S1017 pairing proposed previously is based on a 8/7/9 model [5]. <sup>h</sup> the Pa19-S1017 pairing in the current model (10/6/9).

**Figure 4.**
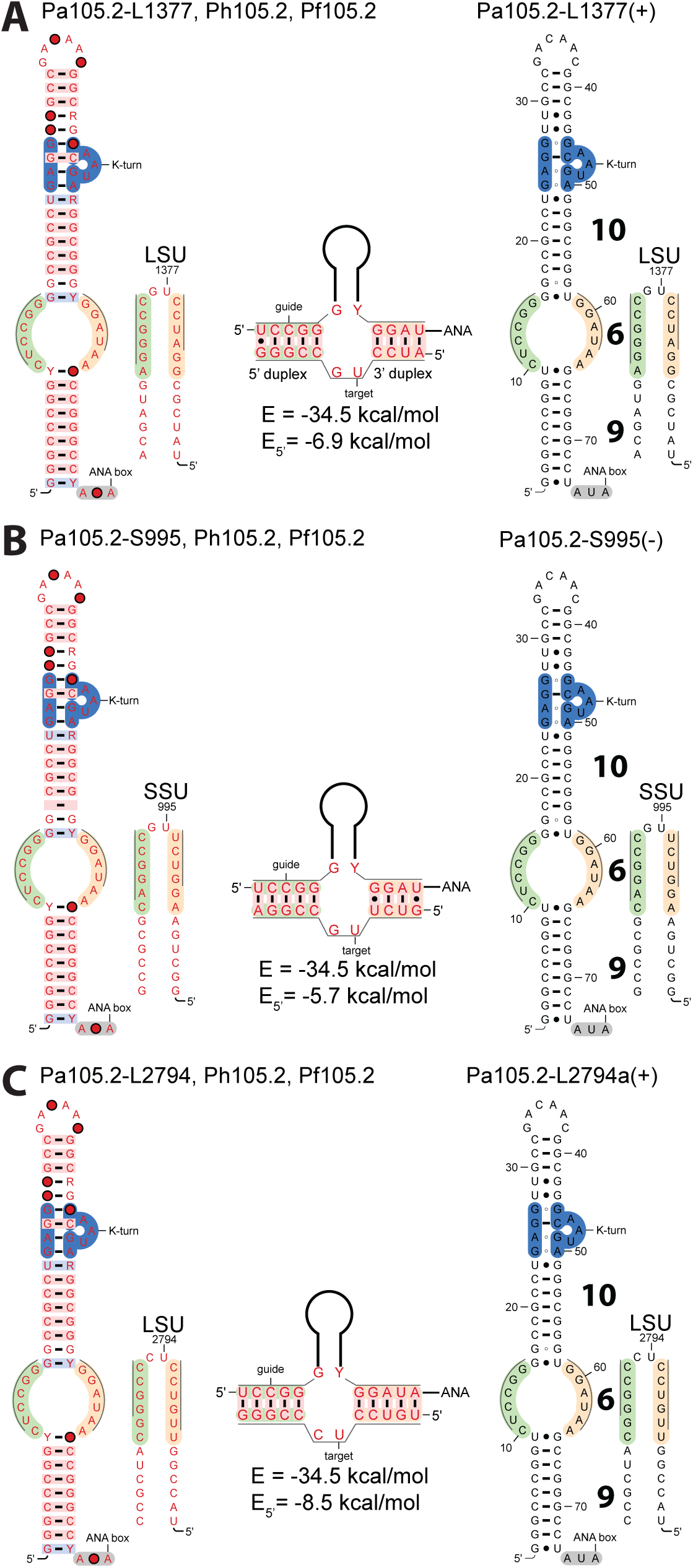
Models of productive (+) and non-productive (−) guide-target pairs of Pa105.2. **A.** Productive fold for Pa105.2 and its true target L1377. **B.** Non-productive fold for Pa105.2 and its false target S995. **C.** Productive fold for Pa105.2 and its true target L2794. The duplexes are represented with the following color code: orange for the 5’ duplex, green for the 3’ duplex. The energy values are those calculated by RNAsnoop.

The structural determinants and the energy criteria proposed in the classification to discriminate between productive/non-productive guide-target pairs are further supported by the analysis of alternative targets identified in the 16S and 23S rRNAs of *P. abyssi* using RIsearch (see Methods). We can define a true target as a ribosomal position which was shown experimentally to be modified by a given H/ACA guide RNA. In the case of *P. abyssi,* there are 16 ribosomal positions corresponding to true targets (pseudo-uridylation: + , Table 1) and 8 other positions corresponding to false targets (pseudo-uridylation: - , Table 1). Among 37 additional false targets (Table S2), 25 guide-pair candidates are expected to be non-productive folds (‘8/7/9’, ‘8/8/9’, ‘9/6/9’, ‘9/6/9’, ‘9/7/8’) while 12 are expected to be productive folds (‘10/5/9’, ‘10/6/9’, ‘10/6/8’). However, all the productive folds except those associated with 3 particular targets (tRNAs) exhibit one of the 7 listed anomalies (Table S2) such as: a 5’ duplex below the energy cutoff (−6 kcal/mol), a mismatch in the 5’ duplex prior to the targeted U position, a succession of wobble pairs in the 5’ duplex, or the absence of pair with the first base of the 3’ duplex (bulge on the guide). The anomalies are expected to have an impact on the stability or on the kinetics of association/dissociation of the guide-pair complex. One particular anomaly corresponds to the presence of a 3 nucleotides spacer between the 5’ and 3’ duplex elements of the target, a feature which was proposed in a previous model for the Pa19-S1017 guide-target complex. This feature is present in two guide-targets pairs (Pa19-L655 and Pa160-L1318, see Table S2) associated with pseudo-uridylation modifications which are not detected in the 23S rRNA of *P. abyssi.* Based on these two examples, this feature is rather considered like an anomaly.

In the particular case of Pa19, the previous model [5] suggested a pairing with only 8 base-pairs in the upper stem and a 3 nucleotide target spacer (Fig. 5A and Supplementary Listing 17): two features which are not compatible with a productive guide-target complex according to our classification. An alternate canonical fold is proposed here corresponding to a ‘10/6/9’ model (Fig. 5B and Supplementary Listing 18) similar to that proposed for Pab105.1-L2554a (Supplementary Fig. S1). The bulge nucleotide is a U which may not stack in the upper stem but rather loop out to the minor groove and still induce a compensatory twist in the upper stem as shown in some known RNA structures [57]. It is thus expected to be productive because of this structural compensation in the upper stem which also includes three stacked G-A base-pairs (Fig. 5). The very long single-strand region at the 5’ side of the internal loop (and the presence of 6 or 7 nucleotides of the 3’ side) may also give enough flexibility to allow the accommodation of the K-turn motif in the proper position and orientation in the RNP particle.

**Figure 5.**
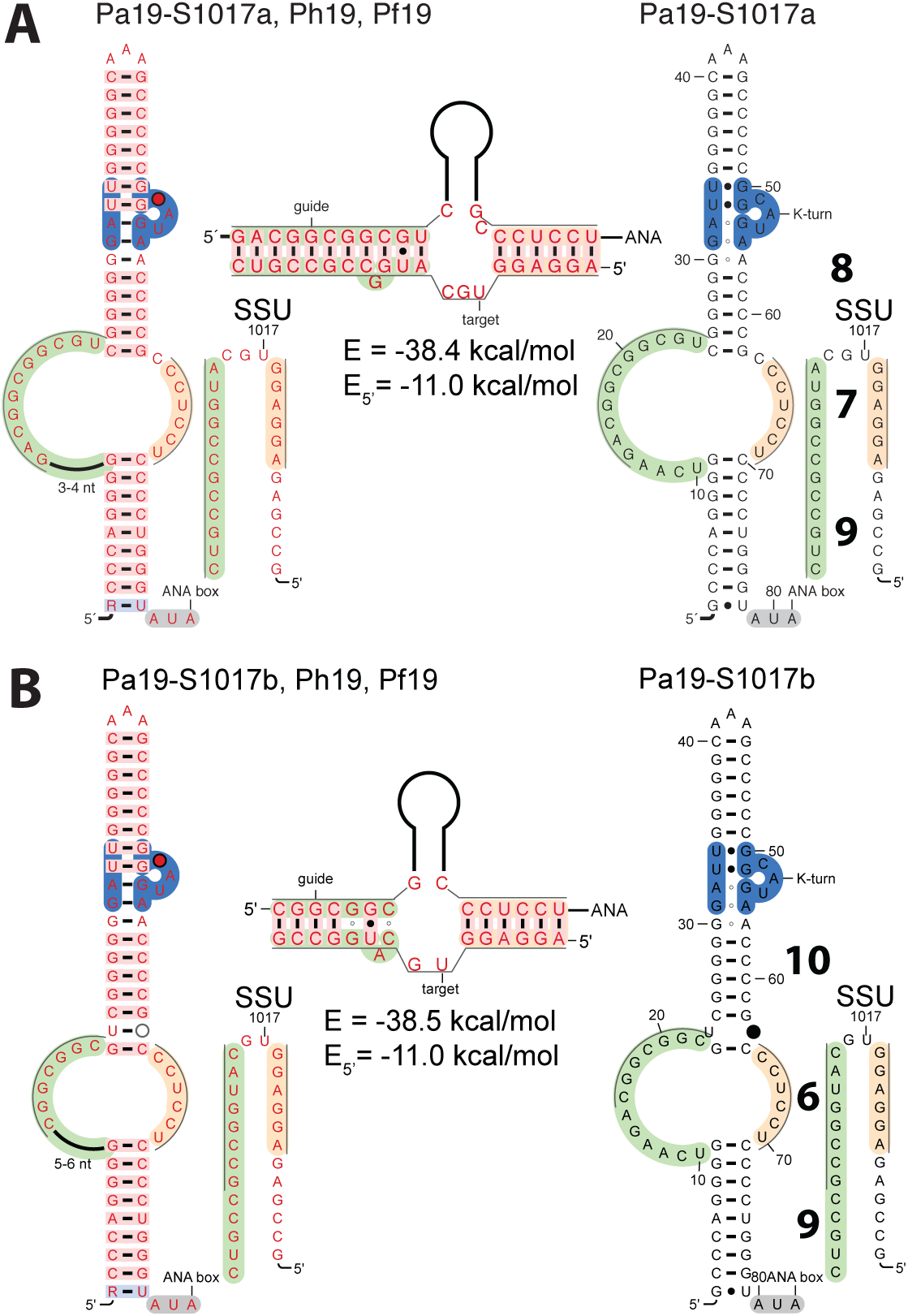
Models of guide-target pairs of Pa19. **A.** Non-productive Pa19-S1017a model proposed previously (‘8/7/9’) [5]. **B.** New productive Pa1 9-S1017b model (‘10/6/9’).

### 2.3 H/ACA RNAs in other Euryarchaea and Crenarchaea

The H/ACA guide RNAs which are present in *Thermococcus* genomes, in particular from *T. kodakaerensis,* are orthologs from *Pyrococcus* motifs as described previously [5]. Only minor sequence variations are observed between *P. abyssi* and *T. kodakaerensis*; the major variation is the substitution in Pa105.1 of the A bulge at the bottom of the upper stem by a Watson-Crick base-pair which restores a more canonical stem (Supplementary Fig. S1). In *Archaeoglobus fulgidus*, all the identified H/ACA guide RNAs fit in the subfamilies described previously in *Pyrococcus*: GUIDE55, GUIDE56, GUIDE65, GUIDE66 and GUIDE75 (Supplementary Fig. S3 & Listing 19). The H/ACA folds are compatible with productive guide RNAs where Af190, Af4.1, Af4.2b and Af46 can associate with their rRNA targets to allow the pseudo-uridylation at the respective positions: S1004, S1167, L2601 and L2639 as shown by Tang *et al.* [1]. It was later suggested that Af52 is not functional for pseudo-uridylation especially because it does not carry any K-turn or K-loop motif [16]. Thus, Af52 is not expected to modify the position L2878 proposed initially [1, 16]. Af4.3 was also proposed to guide the pseudo-uridylation at L1364 but the RNA fold that would target this position (only 9 base-pairs in the upper stem when opening the basal G-U base-pair: see Af4.3 in Supplementary Fig. S3) is not consistent with a productive particle according to our classification. On the other hand, Af4.3 can still guide the modification of L1970 [19]; it is closely related to Pa40.3 and its orthologs in *P. horikoshii* (Ph40.3) and *P. furiosus* (Pf7.3) which target the position L2016. Another related H/ACA motif is also found in *Methanocaldococcus jannaschii* where it can guide the modification at the equivalent position L2015 (Supplementary Fig. S4).

In phylogenetically more distant Euryarchaea such as: *Haloferax volcanii, Haloarcula marismortui, Halobacterium salinarum, Haloquadratum walsbyi* and others, H/ACA RNA guides have also been identified [55, 2]. In the case of *H. volcanii* and the related species mentionned above, the RNA fold proposed based on the association between the guide RNA and its target(s) corresponds to the ‘10/5/9’ fold for the first H/ACA motif (Hvo1 targeting L2621) or the ‘9/7/9’ and ‘11/8/9’ folds for the second motif (Hvo2 targeting L1956 and L1958, respectively) [55]. A slightly modified ‘10/5/9’ fold supported by multiple sequence-structure alignment is proposed for Hvo1 (Supplementary Fig. S5). In the case of Hvo2, a productive fold ‘10/6/9’ can be obtained by extending the upper stem and including an additional unpaired residue downstream of the pseudo-uridylated position as proposed in a previous model of association [5] (Fig. 5); the resulting ‘10/6/9’ fold is annotated Hvo2a (Supplementary Fig. S6). The ‘11/8/9’ alternative fold for Hvo2 includes more than 5 mismatches in the upper stem to accommodate the target sequence [55]. However, a slight change in the pairing of the upper stem and positions of bulge nucleotides both in the guide and target RNAs can make it switch to an alternative ‘10/6/9’ fold suited for the pseudo-uridylation of L1958 (Hvo2b, see Supplementary Fig. S7). Hvo1 and Hvo2a/Hvo2b can be folded according to ‘10/5/9’ and ‘10/6/9’ models to form the following hybrid RNA duplexes: Hvo1-L2621 (Supplementary Fig. S8), Hvo2a-L1956 (Supplementary Fig. S9) and Hvo2b-L1958 (Supplementary Fig. S10).

In the Euryarchaea *M. jannaschii* and *P. furiosus,* differential roles were identified for Cbf5 (TruB/Pus4 family) corresponding to two distinct pseudo-uridylation pathways: a guide RNA-dependent pathway for rRNA modification and a guide RNA-independent pathway for the U55 sequence-specific tRNA modification [58, 25, 29]. In the general case of tRNA modifications, the canonical pseudo-uridylation pathway is based on other Ψ-synthases (PsuX/Pus10 family). In sulfolobales and other Crenarchaea (such as: *Aeropyrum pernix* and *Metalloshaera sedula*), a guide RNA-independent pathway based on an additional Ψ-synthase (TruD/Pus7 family) also operates on tRNAs at position U35 of 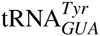. In sulfolobales and related species, this Ψ35-synthase is deficient but rescued through the guide RNA-dependent pathway [20]. The U35 position can be potentially modified through the guide RNA-dependent pathway in 5 species where a compatible H/ACA guide RNA was identified: *Sulfolobus solfataricus, Sulfolobus tokodaii, Sulfolobus acidocaldarius, Aeropyrum pernix, Metallosphaera sedula* [20]. This guide RNA-dependent pathway was validated experimentally for *Sulfolobus solfataricus:* the H/ACA RNA guide Sso1 corresponds to a ‘10/5/9’ fold which is also found in *A. pernix* (Supplementary Fig. S11). On the other hand, the RNA guide candidates found in the other species fold into a ‘9/5/9’ model (Supplementary Fig. S11). However, the tRNA substrates modified through the RNA-dependent pathway have some particularities [58, 20]. Thus, one cannot exclude that the ‘9/5/9’ fold is still active due to compensatory changes associated with the structure of the substrate.

### 2.4 H/ACA(-like) RNAs in Pyrobaculum

From the data provided on the H/ACA-like guide RNAs in *Pyrobaculum* [30], we propose a structural classification which separates the ten sRNA (from Pae sR201 to Pae sR210) into 3 main families which differ by how far they are from the canonical H/ACA motifs found in other archaea, especially in *Pyrococcus.* The first family corresponds to H/ACA motifs reclassified as canonical motifs (Pae sR201 and Pae sR202) while the two other ones correspond to the atypical H/ACA motifs, designated as H/ACA-like motifs. The first two families adopt H/ACA(-like) folds that are consistent with productive complexes according to the structural and functional classification. The third family just includes one of the guide RNA (Pae sR207) and represents a special case that will be discussed later.

The first family includes sR201 and sR202 which can be considered as canonical H/ACA motifs (‘10/5/8’ fold), as shown in Figure 6: the two subfamilies Pae HACA-GUIDE55 and Pae HACA-GUIDE56 are already known from *Pyrococcus* (Fig. 2, Supplementary Listing 20). However, only the sRNA from the Pae HACA-GUIDE55 subfamily (or equivalent) are expected to be productive. All the H/ACA motifs from the Pae HACA-GUIDE55 subfamily exhibit 10 base-pairs between the internal loop and the K-turn motif. On the other hand, the Pae HACA-GUIDE56 subfamily only includes 9 base-pairs, the only two members of this subfamily are from *P. calidifontis* and *P. arsenaticum* (sR202). These two sRNA may switch from the Pae HACA-GUIDE56 to the Pae HACA-GUIDE55 subfamily by shortening the internal loop by one nucleotide which is then included in the upper stem (Supplementary Fig. S12). This leads to increase the number of stacked layers from 9 to 10 to restore a productive configuration (Fig. 6). The minor variation with respect to the canonical H/ACA motif is the length of the lower stem: 8 instead of 9 base-pairs. We can assume that the presence of bulge(s) and/or mismatch(es) in the lower stem allows the guide RNA to be accommodated into the H/ACA RNP particle in a similar way. The ANA box is well conserved in all the motifs sR201 and sR202 from the different species of *Pyrobaculum.* A single variation is observed in *P. calidifontis* where the ANA box is replaced by a CCA box reminiscent to a tRNA 3’ end that might indicate a slight difference in the structural contraints for the guide RNA to be functional.

**Figure 6.**
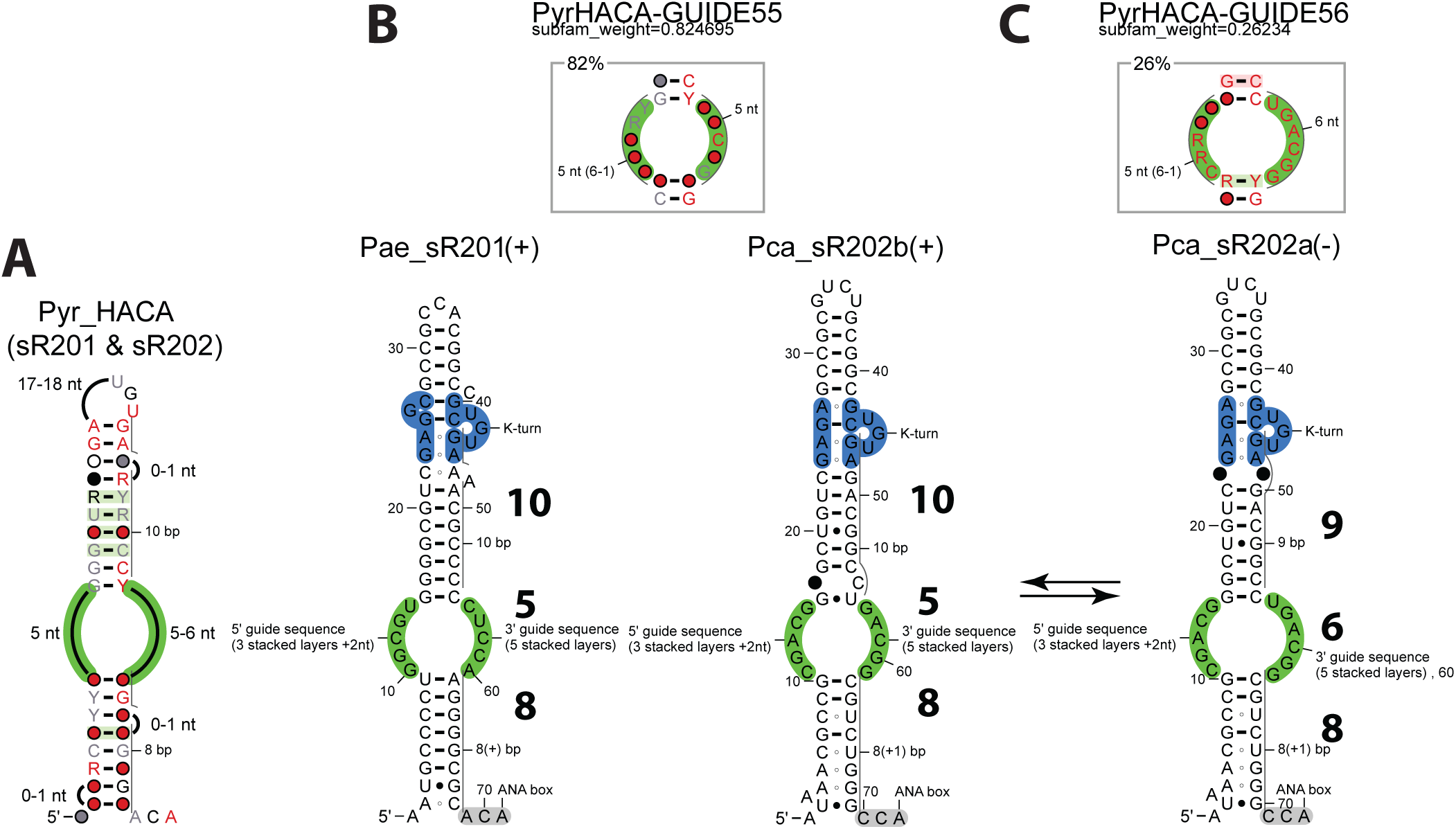
Canonical H/ACA Motifs in *Pyrobaculum* (Pae sR201 & Pae sR202). **A.** Consensus structure of the H/ACA motifs. **B.** Structural subfamily Pae HACA-GUIDE55. **C.** Structural subfamily Pae HACA-GUIDE56.

The second family includes all the other H/ACA-like guide RNAs (sR203 to sR210) except sR207. Although they are truncated from the lower stem, they generally include one or two possible residual base-pair(s) at the position(s) in the sequence which would be consistent with the Pae HACAGUIDE66 or Pyr HACA-GUIDE65 subfamilies (Fig. 7A). From the structural alignment of the H/ACA-like motifs for this family (Supplementary Listing 21): 64% of the analyzed sequences do contain one or two base-pairing position(s) (subfamiles BPS, BP1 and BP2, Fig. 7B-D) reminiscent from the canonical H/ACA motif while 36% lost any trace of the lower stem (subfamily NONE, Fig. 7E). Representative members of each subfamily are shown: the presence of residual base-pairs in about two thirds of the H/ACA-like motifs suggest they may derive from canonical H/ACA motifs degenerated by the accumulation of mutations in the lower stem. The H/ACA-like motifs generally carry an ANA box or a more degenerated NCA box at a distance from the stem which is consistent with the Pyr HACA-GUIDE66 or Pyr HACA-GUIDE65 subfamilies (modulo one residue) and the canonical H/ACA motifs (59%). Some motifs (27%) still carry a degenerated box (NCA) which is downstream of the expected position (up to 10-12 residues). In a few cases (14%), there is no ANA or NCA box (Supplementary Fig. S13).

**Figure 7.**
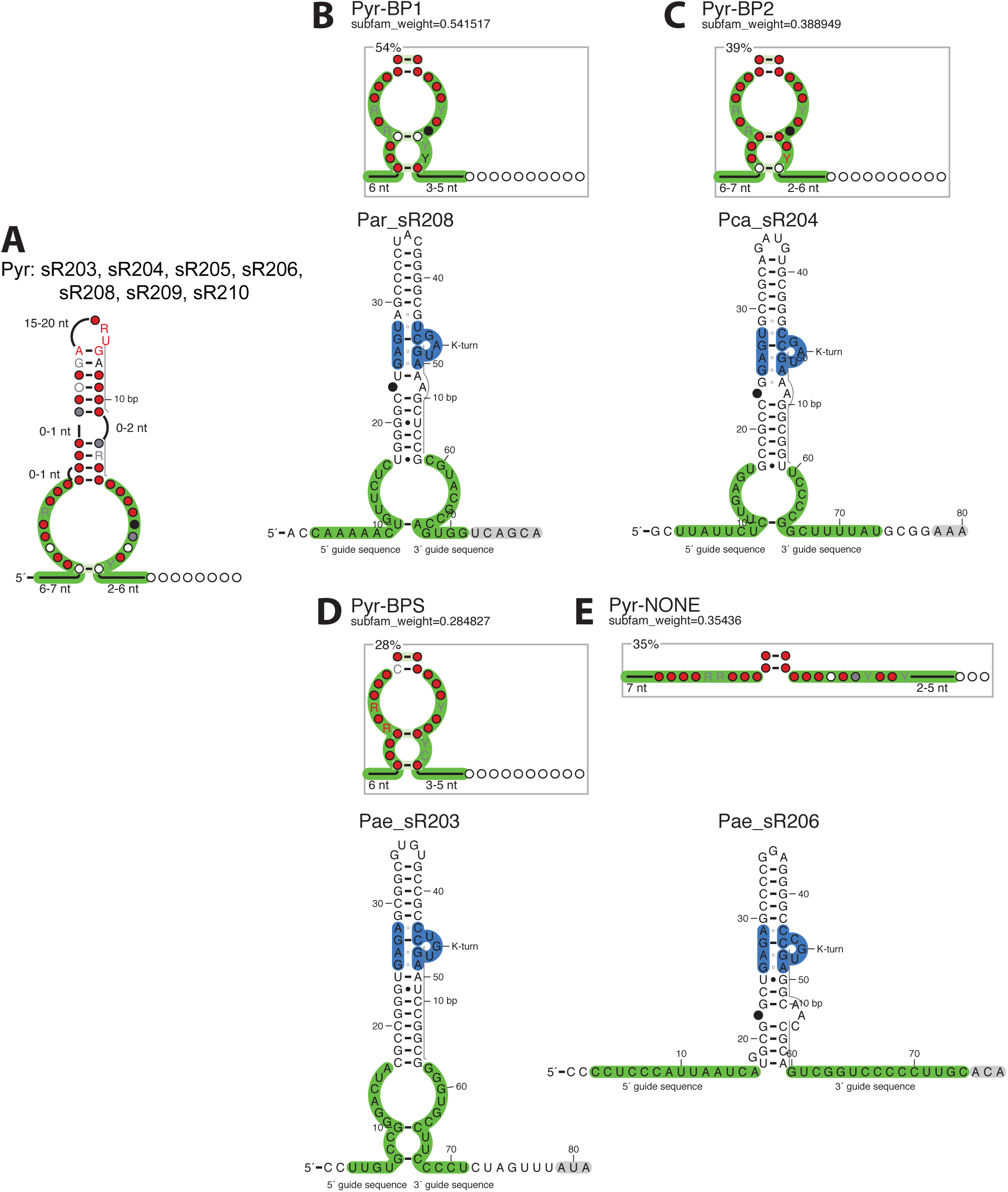
H/ACA-like Motifs in *Pyrobaculum.* **A.** Consensus structure of all the H/ACA-like motifs. **B.** Structural subfamily (Pyr-BP1) with one pseudo base-pair corresponding to the (n+6) position in the canonical lower stem. **C.** Structural subfamily (Pyr-BP2) with one pseudo base-pair corresponding to the (n+9) position in the canonical lower stem. **D.** Structural subfamily (Pyr-BPS) with two pseudo base-pairs corresponding to the (n+6) and (n+9) positions in the canonical lower stem. **E.** Structural subfamily (Pyr-NONE) without any pseudo base-pair.

The associations between Pae sR207 and its two ribosomal targets L2597 and L2596 [30] are based on a 12/(13)- and 13/(12)- model of pairing, respectively. There is no direct evidence that these two ribosomal positions are actually modified by Pae sR207 and none of the paring models is expected to be productive (Table 2 and Fig. 8). The L2596 position is particularly questionable since it requires the presence of two successive wobble base-pairs closing the pseudo-upper stem while the presence of a second U-G base-pair is not supported by phylogenetic data (Supplementary Fig. S14 and S15). Alternative folds that would restore a productive H/ACA-like motif can be proposed by excluding some nucleotides from the stacking layers of the stem (Fig. 9, Supplementary Listing 22). However, there is no compatible compensation that would allow Pae sR207 to target L2596. Alternative targets were identified in 16S, 23S rRNAs and tRNAs (Table S3) but half of the guide-target pairs exhibit a weak 5’ duplex (below the energy cutoff of −6 kcal/mol) and the other half some pairing anomaly(ies) similar to those already described in *P. abyssi.*

**Table 2.**
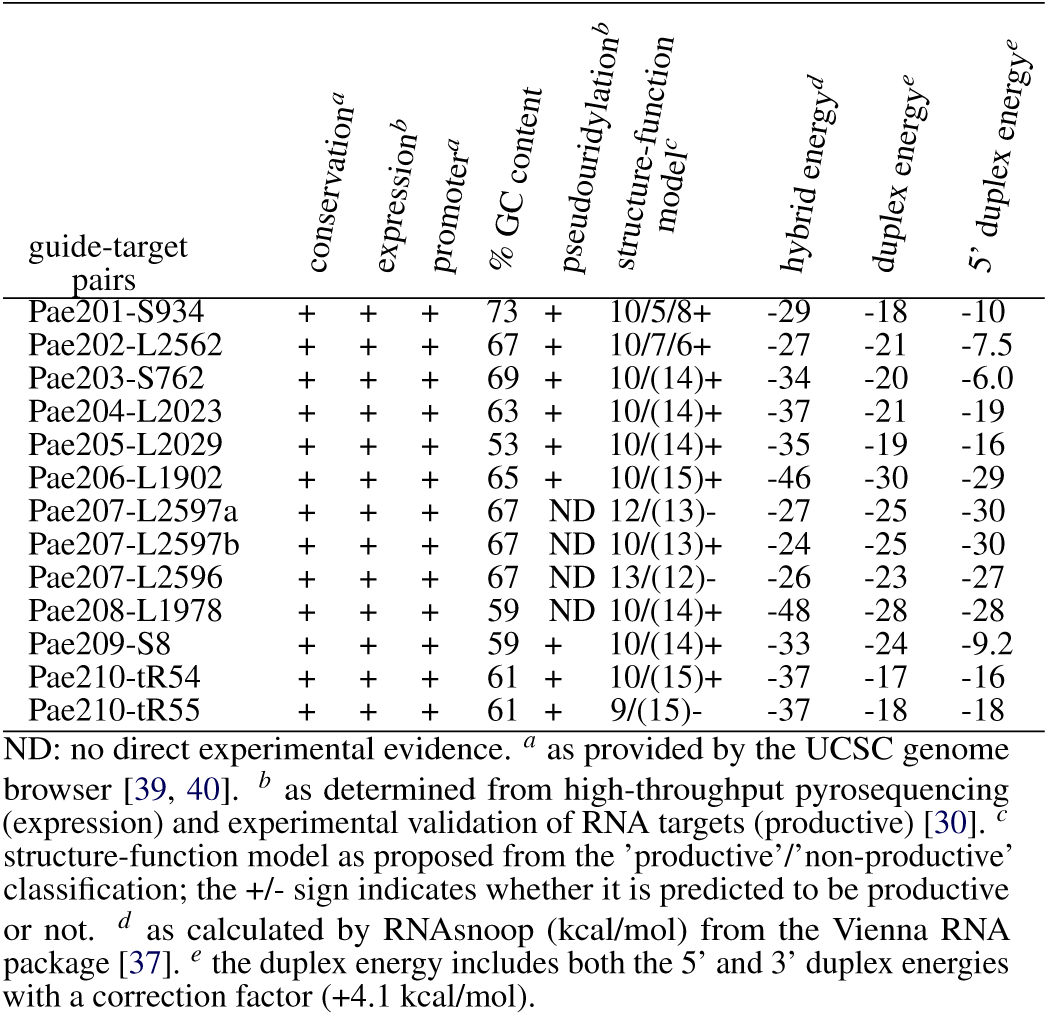
Functional and structural features from known H/ACA guide RNAs in *Pyrobaculum* targeting rRNAs and tRNAs.

| guide-target pairs | conservation<sup>a</sup> | expression<sup>b</sup> | promoter<sup>a</sup> | % GC content | pseudouridylation<sup>b</sup> | structure-function model<sup>c</sup> | hybrid energy<sup>d</sup> | duplex energy<sup>e</sup> | 5' duplex energy<sup>e</sup> |
| --- | --- | --- | --- | --- | --- | --- | --- | --- | --- |
| Pae201-S934 | + | + | + | 73 | + | 10/5/8+ | -29 | -18 | -10 |
| Pae202-L2562 | + | + | + | 67 | + | 10/7/6+ | -27 | -21 | -7.5 |
| Pae203-S762 | + | + | + | 69 | + | 10/(14)+ | -34 | -20 | -6.0 |
| Pae204-L2023 | + | + | + | 63 | + | 10/(14)+ | -37 | -21 | -19 |
| Pae205-L2029 | + | + | + | 53 | + | 10/(14)+ | -35 | -19 | -16 |
| Pae206-L1902 | + | + | + | 65 | + | 10/(15)+ | -46 | -30 | -29 |
| Pae207-L2597a | + | + | + | 67 | ND | 12/(13)- | -27 | -25 | -30 |
| Pae207-L2597b | + | + | + | 67 | ND | 10/(13)+ | -24 | -25 | -30 |
| Pae207-L2596 | + | + | + | 67 | ND | 13/(12)- | -26 | -23 | -27 |
| Pae208-L1978 | + | + | + | 59 | ND | 10/(14)+ | -48 | -28 | -28 |
| Pae209-S8 | + | + | + | 59 | + | 10/(14)+ | -33 | -24 | -9.2 |
| Pae210-tR54 | + | + | + | 61 | + | 10/(15)+ | -37 | -17 | -16 |
| Pae210-tR55 | + | + | + | 61 | + | 9/(15)- | -37 | -18 | -18 |

ND: no direct experimental evidence. <sup>a</sup> as provided by the UCSC genome browser [39, 40]. <sup>b</sup> as determined from high-throughput pyrosequencing (expression) and experimental validation of RNA targets (productive) [30]. <sup>c</sup> structure-function model as proposed from the 'productive'/'non-productive' classification; the +/- sign indicates whether it is predicted to be productive or not. <sup>d</sup> as calculated by RNAsnoop (kcal/mol) from the Vienna RNA package [37]. <sup>e</sup> the duplex energy includes both the 5' and 3' duplex energies with a correction factor (+4.1 kcal/mol).

**Figure 8.**
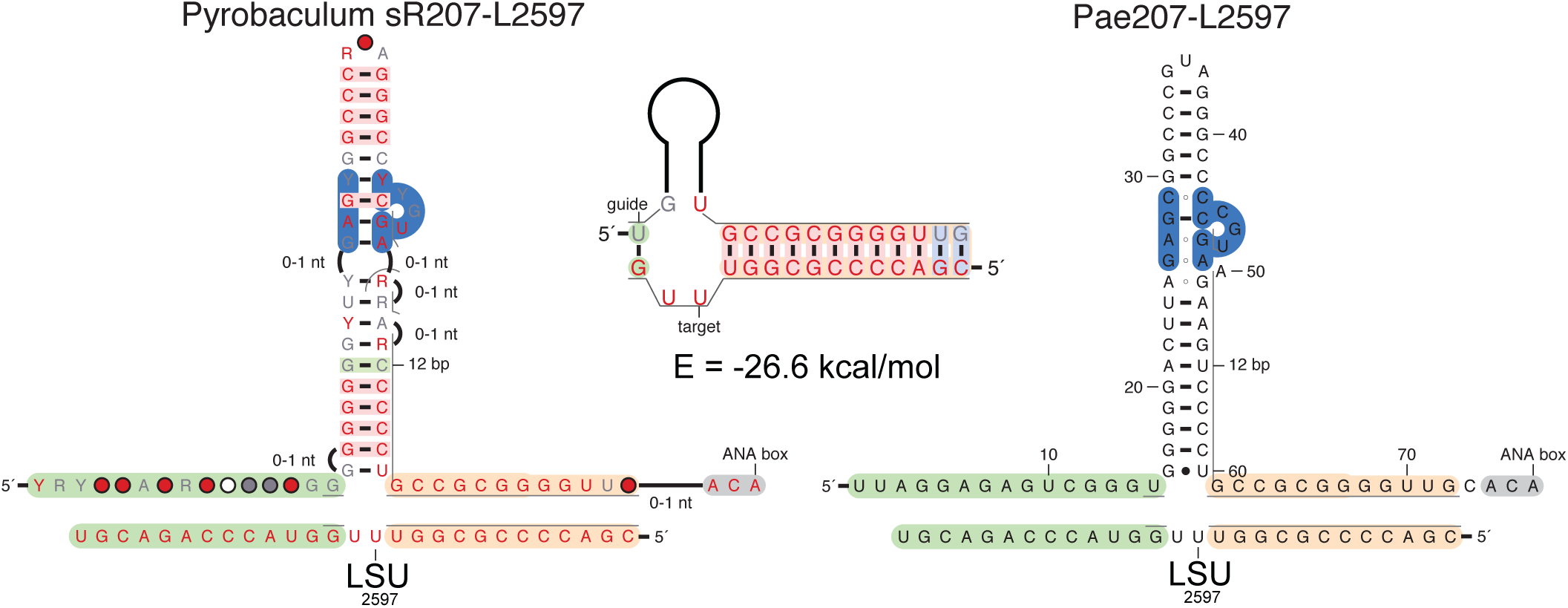
Pairing Model of guide-target Pae sR207-L2597

**Figure 9.**
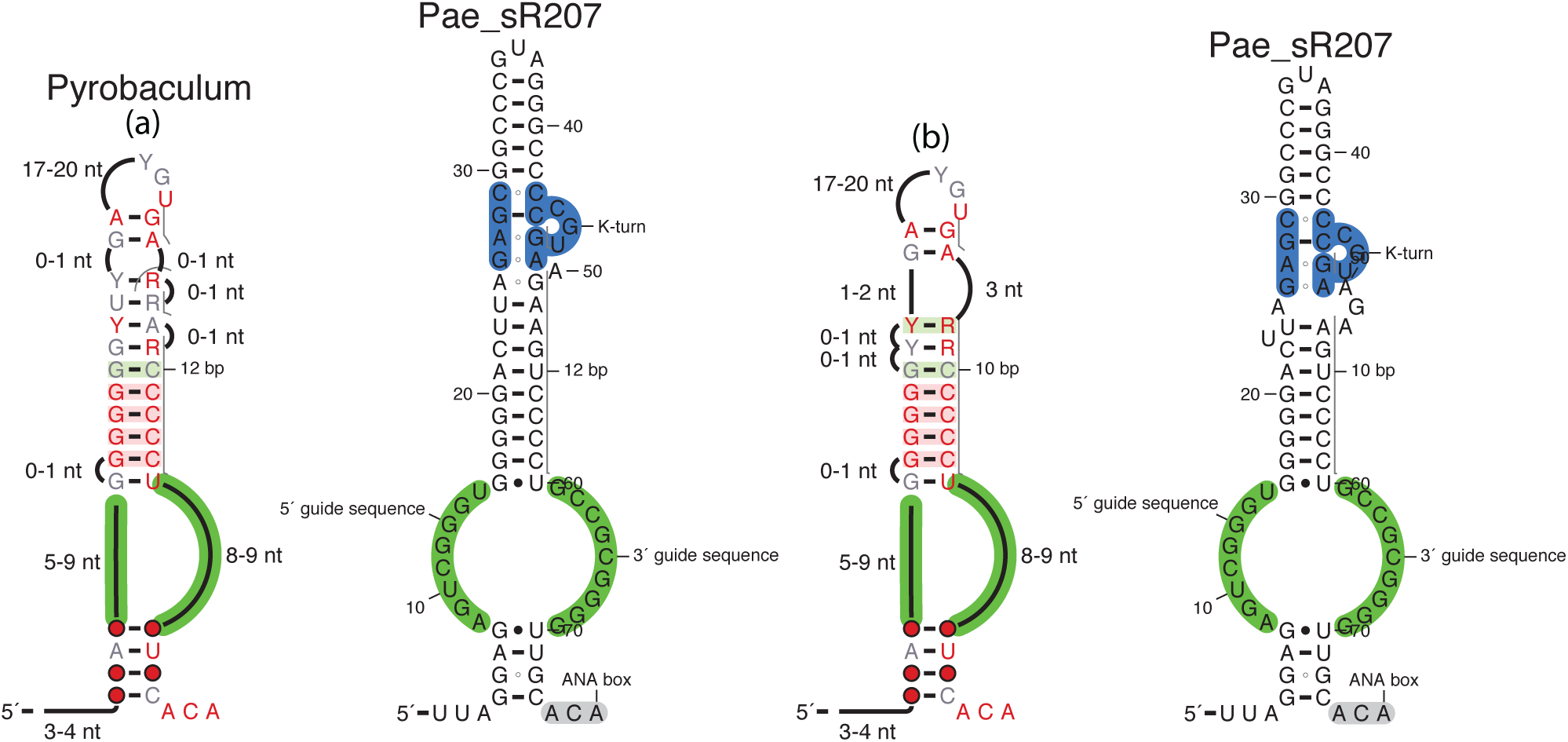
Alternative RNA folds for Pae sR207. (a) RNA fold proposed by Bernick *et al.* [30] vorresponding to a 12/13 model; (b) RNA fold compatible with a 10/(13) model.

### 2.5 New H/ACA(-like) guide candidates in Pyrococcus

Based on the H/ACA-like motifs present in *Pyrobaculum* genomes, the genome of *P. abyssi* was scanned for the presence of similar motifs in transcribed regions [33] (see Fig. 1). Several new H/ACA(-like) motifs were identified and ranked according to several criteria: GC-content, conservation, expression, promoter strength [33]. Among the 8 new candidates thus identified, three of them look like canonical H/ACA motifs with minor variations and the five other ones correspond to H/ACA-like motifs where the lower stem is absent or mostly truncated (Fig. 10). Three of the motifs were initially considered of particular interest because they satisfy at least two of the criteria listed above: PabO1 (high expression, strong promoter, conservation, medium GC-content, Supplementary Fig. S16), PabO48 (high conservation, high expression, Supplementary Fig. S17), PabO78 (high expression, strong promoter, conservation: Supplementary Fig. S18). PabO48 looks like a canonical H/ACA motif while PabO1 and PabO78 are H/ACA-like motifs suggesting these non-canonical motifs might not be specific to Crenarchaea. However, both PabO1 or PabO78 adopt a non-productive fold (’9/10/9’ and ’9/14’, respectively); the ANA box signature is present in both motifs but it is not located at the proper position in PabO1. There are 3 tRNA genes with 12- or 13-mer sequences that could be targeted by PabO1 (tRNA-Pro*^CGG^*, tRNA-Pro*^TGG^*, tRNAAsn*^TAA^*, data not shown) but missing a U at the proper position. PabO48 is the only motif that could target a ribosomal position (L2047) but it does not carry any ANA box signature at the expected position.

**Figure 10.**
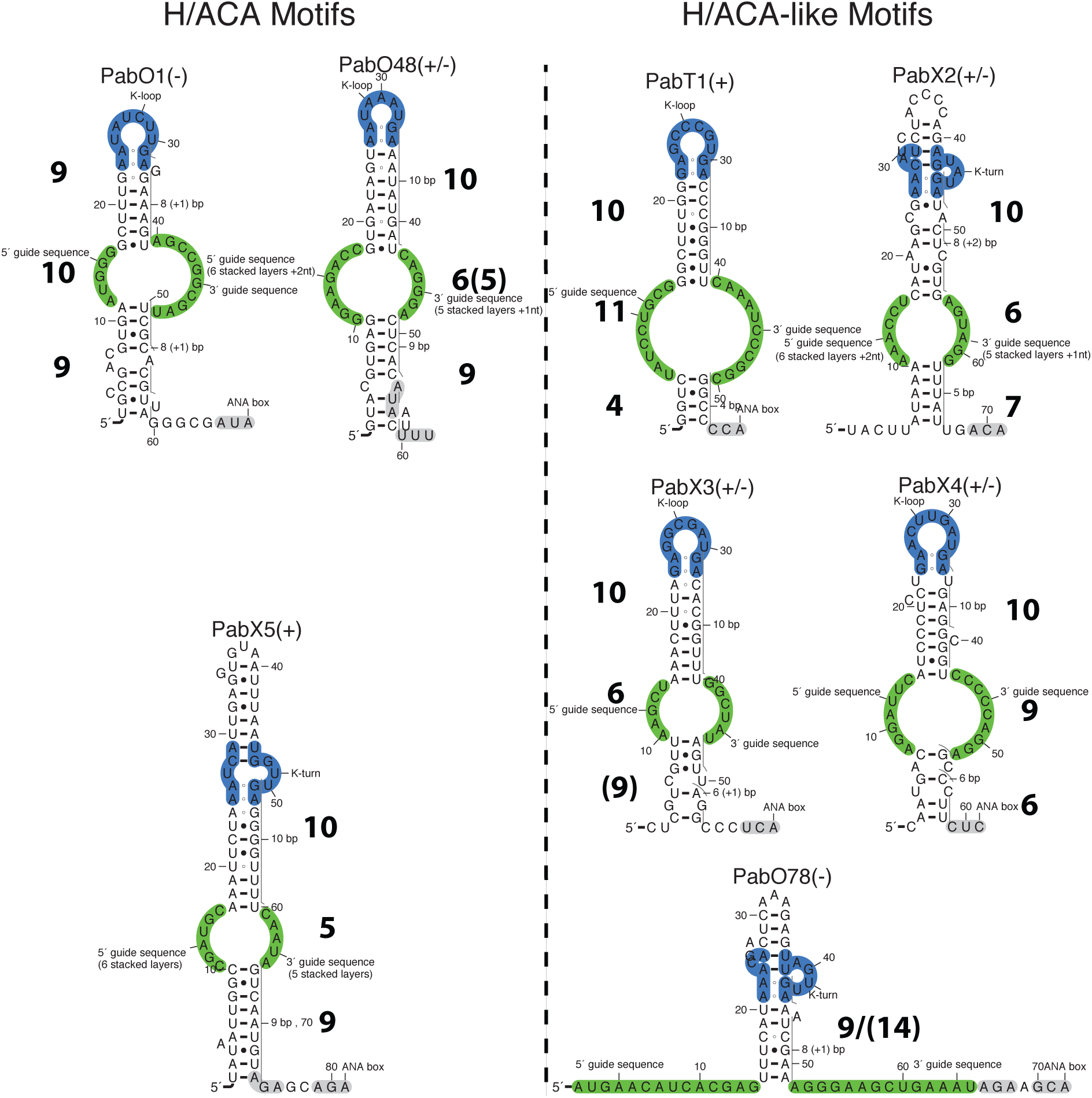
New H/ACA(-like) motifs identified in *Pyrococcus abyssi* [33]. The functional annotation (+, − or +/−) indicates the relevance of the H/ACA(-like) candidate based on the following criteria: (1) productive fold (‘10/6/9’, ‘10/5/9’ or equivalent folds), (2) distance of 14 to 15 residues from the internal loop to the ANA box, (3) ANA box, (4) absence of alternative folds due to pairings in the internal loop. (+): all4 criteria are met; (+/−) only 3 criteria are met; (−) less than 3 criteria are met.

A fourth motif, PabT1 (Supplementary Fig. S19), also meets the indicated criteria but it is embedded within the 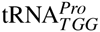 gene and no potential target was found in the whole genome for this motif. The other 4 motifs are annotated PabX2 to PabX5 (Supplementary Fig. S20 to Fig. S23): they are all conserved in at least two *Pyrococcus* species and located in expressed regions except PabX2. Only PabX4 has a strong promoter and only PabX5 corresponds to a canonical H/ACA motif. PabX5 is the only canonical motif that includes a proper ANA box but there is no potential targeted sequence in the genome. PabX2 and PabX3 can adopt alternative folds involving several pairings in the internal loop suggesting they may be degenerated H/ACA-like motifs or just false-positive candidates.

### 2.6 Extra-Ribosomal Targets from H/ACA(-like) RNAs in Pyrococcus and Pyrobaculum

The full genomes of *P. abyssi* and *P. aerophilum* were explored to identify extra-ribosomal targets associated with H/ACA motifs from *P. abyssi* and H/ACA(-like) motifs from *P. aerophilum* (see Materials and Methods). A large number of potential targets were identified since the sequence constraints for guide-target pairing are rather loose. In *Pyrococcus abyssi,* more than 4000 hits were obtained: only 37% of those correspond to productive folds (‘10/5/9’ or ‘10/6/9’ or equivalent folds), the remaining 63% correspond to non-productive alternative folds for all the H/ACA motifs. Hits from both the productive and non-productive folds are more frequently found in 16S and 23S rRNAs that represent 0.3% of the genome size. The hits from the productive motifs have a 10-fold to a 3-fold enrichment (with respect to the expected number of targets by chance in rRNA) whether one considers or not the overlapping targets (Fig. S24). On the contrary, there is an under-representation of targets in CDS regions (92% of the genome size) due to a similar proportion of targets (around 50%) located in antisense from CDS or in other functional regions. In *P. aerophilum,* more than 6,000 hits were obtained: hits from both the productive and non-productive folds are only slightly more frequent in 16S and 23S rRNAs (2 to 5- fold over-representation). In hits from productive folds, the targets located in 16S and 23S rRNAs represent 0.3% of the non-overlapping targets while these regions of the genome represent 0.2% of the genome size (Fig. S25). The number of targets in the CDS regions is much more under-represented due to a large number of targets in antisense from CDS or other functional regions.

There is a total of 525 common CDS between *P. abyssi* and *P. aerophilum* but only 455 correspond to annotated genes (after excluding hypothetical proteins). Among the 525 CDS, there are 246 CDS targeted by H/ACA(-like) motifs from both *P. abyssi* and *P. aerophilum* (Fig. 11); among the 246 CDS, 218 correspond to annotated genes.

**Figure 11.**
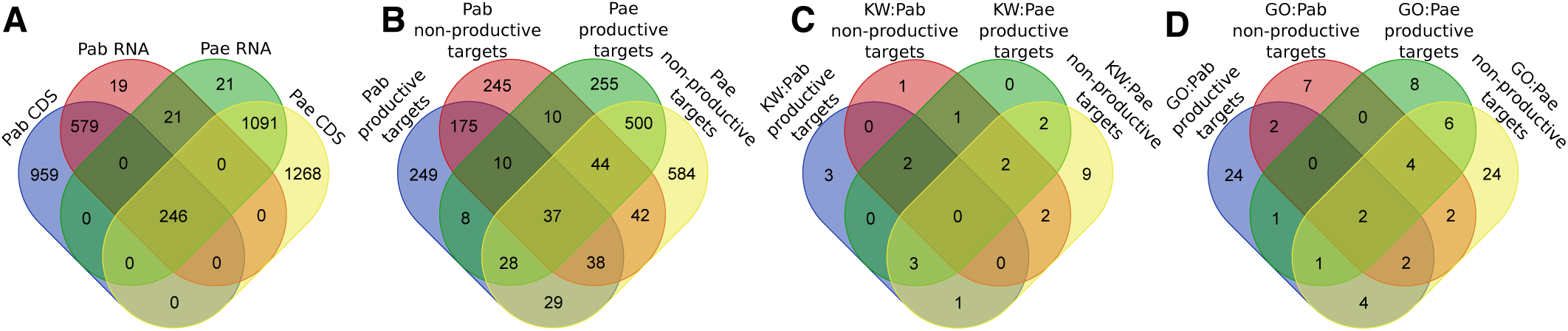
Venn Diagrams of CDS and Associated Functions Targeted by H/ACA(-like) folds from *P. abyssi* (Pab) and *P. aerophilum* (Pae). **A.** CDS genes/RNA genes targeted by Pab and Pae (full list of genes: Listing 23). **B.** CDS targeted by productive/non-productive H/ACA(-like) motifs from Pab and Pae (full list of CDS: Listing 24). **C.** Biological functions or processes (UniProt keywords) targeted by productive/non-productive H/ACA(-like) folds from Pab and Pae (full list of keywords: Listing 25). **D.** Biological functions or processes (GO terms) targeted by productive/non-productive H/ACA(-like) folds from Pab and Pae (full list of GO terms: Listing 26).

In the sublist of 246 genes which are potential common targets in both *Pyrococcus* and *Pyrobaculum* (Fig. 11A-B), the more statistically relevant functions (DAVID analysis with P value lower than 1E-4 and Benjamini score lower than 1E-3, Fig. 11C-D) are: the amino-acid biosynthesis and nitrogen compound biosynthetic process (UniProt: KW-0028 & GO:0044271, enrichment score of 4.09), zinc (UniProt: KW-0862, enrichment score of 2.68), nucleotide-binding and ATP-binding (UniProt: KW-0547 and KW-0067, enrichment score of 2.33). Interestingly, the gene of the 3-dehydroquinate synthase (ENTREZ Gene ID: 1495344) found in the cluster of amino-acid biosynthesis was identified as potentially regulated by some guide RNA (C/D box or H/ACA-like) in *Pyrobaculum calidifontis* [59]. Similarly, one gene corresponding to the elongation factor EF-2 (ENTREZ Gene ID: 1495166) and two genes corresponding to tRNA-synthetases are potentially regulated by guide RNAs in different species of *Pyrobaculum* [59]. Among the genes candidates clustered in the nucleotide-binding function, we also find EF-2 and 9 tRNA-synthetases which makes this latter class of genes the more abundant category among all gene candidates and perhaps the more relevant candidates to test.

In the extra-ribosomal targets, the guide-target hybrid duplexes generally correspond to sub-optimal hits including more wobble pairs and/or bulge(s) and mismatch(es); the more frequently involved H/ACA(-like) RNAs are: Pab19 for *P. abyssi* and Pae sR207 for *P. aerophilum.* As an example, the leucyl-tRNA synthetase (ENTREZ Gene ID: 1496220) is shown as a potential gene target (Supplementary data: Fig. S26). As proposed previously in *Pyrobaculum* [59], some of the H/ACA(-like) RNAs may act as antisense RNAs but it is not clear whether the full-length RNA is involved [59]. The data provided here suggest the H/ACA(-like) RNAs might potentially act as specific antisenses taking advantage of the H/ACA RNP machinery to stabilize some productive or nonproductive complexes with mRNA targets.

## 3. Conclusions

A more precise structural and functional classification of archaeal H/ACA guide RNAs is proposed based on the X-ray data of H/ACA RNPs and 2D RNA predictions supported by the phylogenetic data of the known H/ACA(-like) motifs and their target(s); it is consistent with all the current data available up to now in both Euryarchaea and Crenarchaea. One of the major structural determinants is the presence of a stretch of 10 stacked layers in the upper stem from the closing base-pair of the pseudo-uridylation pocket to the G-A sheared base-pairs of the K-turn or K-loop motif. A distance of 14 or 15 nucleotides from the same closing base-pair to the ANA box proposed previously as another structural determinant [5] is the result of the conformational flexibility at the junction between the internal loop and the lower stem. The 3’ single-stranded region of the internal loop usually varies between 5 and 6 nucleotides in the typical RNA folds: in the GUIDE65 subfamily (5 nucleotides) the helical twist is more pronounced at the junction than in the GUIDE66 subfamily (6 nucleotides) and compensates for the shorter 3’ single-stranded region. Finally, the stability of the 5’ duplex also appears to play a key role in the formation of productive RNA-RNA complexes. It probably contributes to tether the uridine to be modified in the pseudo-uridylation pocket and to restrain its mobility in the catalytic site of Cbf5.

A more detailed structural analysis of H/ACA-like guide RNAs in the *Pyrobaculum* genus reveals a strong similarity with the canonical H/ACA guide RNAs previously identified in both Euryarchaea and Crenarchaea. Two particular sRNAs (sR201 and sR202) can actually be considered as canonical H/ACA guide RNAs. The other H/ACA-like guide RNAs (sR203 to sR210) can be classified in different subfamilies which retain to some degree the vestiges of the lower stem. The H/ACA(-like) motifs identified recently in *P. abyssi* [33] show a similar signature with a pseudo lower stem degenerated by the incorporation of mismatches and bulges. According to this current work, all the H/ACA(-like) motifs from *P. aerophilum* except one are classified as productive as expected from the published data [30]. In the particular case of Pae sR207, its role in the modification of the positions L2596 and L2597 is not confirmed because there is no direct experimental evidence; an alternative RNA fold that is consistent with a 10/(13) model is proposed for the modification of L2597. The presence of genes which might be regulated by H/ACA(-like) RNAs, a hypothesis which was already proposed based on experimental evidence in *Pyrobaculum* [59], remains to be tested in the case of gene candidates which are potential targets in both *Pyrococcus* and *Pyrobaculum.*

## Acknowledgments

The bioinformatics workflow integrates S-MART scripts which were developed by Matthias Zytnicki. Perl scripts were also provided by Olivier Lespinet. The authors are members of the french network “GDR-Archaea”.

